# An ABA-GA bistable switch can account for natural variation in the variability of Arabidopsis seed germination time

**DOI:** 10.1101/2020.06.05.135681

**Authors:** Katie Abley, Pau Formosa-Jordan, Hugo Tavares, Emily Chan, Ottoline Leyser, James C.W. Locke

**Affiliations:** The Sainsbury Laboratory, University of Cambridge, Cambridge, Cambridgeshire, United Kingdom, CB2 1LR

**Author notes:** Corresponding authors: (JCWL) (OL). These authors contributed equally (Katie Abley, Pau Formosa-Jordan).

## Abstract

Genetically identical plants growing in the same conditions can display heterogeneous phenotypes. Whether this phenotypic variability is functional and the mechanisms behind it are unclear. Here we use Arabidopsis seed germination time as a model system to examine phenotypic variability. We show extensive variation in seed germination time variability between Arabidopsis accessions, and use a multi-parent recombinant inbred population to identify two loci involved in this trait. Both loci include genes implicated in ABA signalling that could contribute to seed germination variability. Modelling reveals that the GA/ABA bistable switch underlying germination can amplify variability and account for the effects of these two loci on germination distributions. The model predicts the effects of modulating ABA and GA levels, which we validate genetically and by exogenous addition of hormones. We confirm that germination variability could act as a bet hedging strategy, by allowing a fraction of seeds to survive lethal stress.

## Introduction

In an environment where current cues cannot be used to predict future conditions, a bet-hedging strategy can be advantageous, whereby the same genotype produces a variety of phenotypes in individuals in a common environment (Cohen, 1966; Lewontin and Cohen, 1969; Simons, 2011). Bacteria can use bet-hedging strategies to survive antibiotic treatments (Balaban et al., 2004; Martins and Locke, 2015). Mechanistic studies suggest that the required phenotypic variability is generated by genetic networks amplifying stochasticity in molecular interactions to generate a range of outputs (Alon, 2007; Eldar and Elowitz, 2010; Viney and Reece, 2013).

In plants, theoretical work shows that variability in seed germination time is likely to be advantageous in environments that are unpredictable (Cohen, 1966; Simons, 2011). Indeed, ecological studies have found that variability in seed germination time is correlated with environmental unpredictability (Simons and Johnston, 2006; Venable, 2007). This variability can involve germination of genetically identical seeds being spread between seasons or within a season (or a combination of both), and all these behaviours exist in wild species.

The mechanisms underlying how seed germination is spread between seasons have been studied in detail. Seeds can enter a ‘dormant’ state, refractory to germination even under favourable germination conditions (Baskin and Baskin, 2004; Bewley, 1997). Seed dormancy is a continuous variable, with genetically identical seeds having different depths of dormancy (Finch-Savage and Leubner-Metzger, 2006). However, the extent of dormancy for a batch of seeds is usually estimated by quantifying the percentage germination, which does not provide information about the distribution of germination times of individual seeds that germinate within a season, or experiment.

Although variability in germination times within a season has been studied in desert annuals (Simons and Johnston, 2006), these species are not amenable to genetic or mechanistic studies. Little is known about variability in germination time within a season in the model plant *Arabidopsis thaliana*. The extent of variability, the mechanisms that underlie it, or how related the underlying mechanisms are to those that control seed dormancy between seasons, are unknown. However, there is a large body of work using percent germination as a measure of seed dormancy. A number of quantitative genetic studies have identified loci that underlie natural variation in the extent of dormancy under different environmental conditions, with the *DELAY OF GERMINATION* (*DOG*) loci being the first identified and some of the most characterised (Alonso-Blanco et al., 2003; Bentsink et al., 2010; Clerkx et al., 2004; Footitt et al., 2019; Kerdaffrec and Nordborg, 2017; Meng et al., 2008; van Der Schaar et al., 1997). The molecular mechanisms underlying germination have also been uncovered. In-depth molecular studies have shown that the decision to germinate is controlled by the balance between two hormones, gibberellic acid (GA), which promotes germination, and abscisic acid (ABA), which represses it (Liu and Hou, 2018). These hormones function in a mutually antagonistic manner by each inhibiting the synthesis and promoting the degradation of the other (Liu and Hou, 2018; Piskurewicz et al., 2008). Additionally, the two hormones have opposing effects on downstream transcriptional regulators that control the balance between dormancy and germination (Liu et al., 2016; Piskurewicz et al., 2008; Shu et al., 2013). Pioneering modelling work has suggested that variable germination times can be generated by variation in sensitivities to germination regulators in a batch of seeds (Bradford, 1990; Bradford and Trewavas, 1994), or due to stochastic fluctuations in the regulators (Johnston and Bassel, 2018). However, it remains unclear how different genotypes generate different extents of variability in germination times. Additionally, the extent to which there is natural variation in germination time distributions in Arabidopsis has not been described.

Here we set out to investigate germination time variability in Arabidopsis. We demonstrate that there is robust natural variation in germination time variability, and that this is genetically separable from mode germination time (i.e. the timing of the peak of the distribution), allowing these properties to be tuned independently. We show that two loci underlie variability in germination time. One of these loci overlaps with the *DOG6* locus and its effect on variability in germination time is correlated with effects on the mode, while the other overlaps with the *DOG1* locus and affects variability independently from the mode. We show that variability in germination time appears to be an inherent property of each seed, rather than being generated by seed position on the parent plant. Based on these findings, we generate a mathematical model of the GA/ABA bistable switch that underlies germination. We show that the switch can amplify variability and account for the differing effects of the two identified loci on different germination traits. We validate the model by testing predictions about the effects of perturbations to ABA and GA levels. Finally, we show that increased variability in germination time can act as a bet hedging strategy against short durations of lethal stresses, suggesting circumstances under which selection for variability could confer a selective advantage.

## Results

### Variability in seed germination time varies genetically in Arabidopsis

We first determined whether Arabidopsis exhibits natural variation in the variability of seed germination time. To do this, we quantified germination time distributions for 19 natural accessions and a multi-parent recombinant inbred line population (MAGIC population), derived from those accessions (Kover et al., 2009). We also included 10 lines that we selected from a set of Spanish accessions as being likely to have low or high variability in germination time based on their germination time distributions over the first 6 days after sowing (Vidigal et al., 2016). We grew plants in controlled conditions for seed harvesting, and collected all the seeds from 3 plants of each line. After a fixed period of dry storage (∼30 days), we sowed a sample of each of these replicate batches of seeds in petri-dishes in controlled conditions and scored germination every day until there had been no further germination for a period of 2 weeks (see methods for further details). We chose a relatively short period of dry storage for our experiments as we reasoned that this would best reflect the state of naturally shed seeds. In these conditions, the MAGIC lines had low levels of seed dormancy (with 30 days of dry storage, 82% of lines had >= 50% germination), allowing us to quantify germination time distributions and estimate its variability. We used the coefficient of variation (CV =standard deviation / mean) of the germination time distribution as a measure of variability. We confirmed that CVs for the MAGIC parental lines remained similar over a range of lengths of dry storage period (30-60 days of dry storage), demonstrating that our results are not specific to one condition (Figure 1-figure supplement 1A, B).

**Figure 1.**
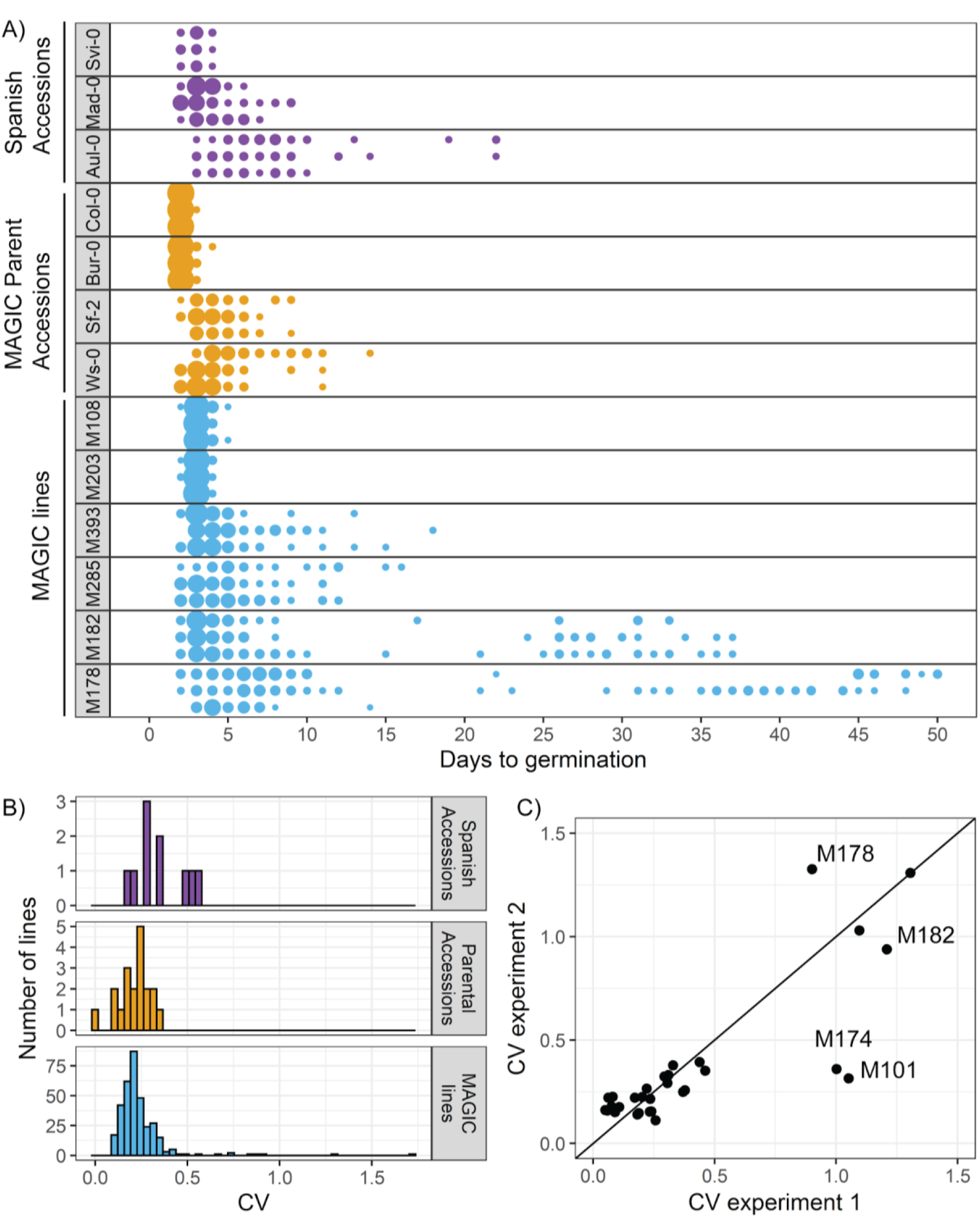
There is variation in variability in germination times in Arabidopsis. **A)** Examples of distributions of germination time for natural accessions and MAGIC lines. Each row shows the germination time distribution of a seed batch from a different parent plant of a particular line, colours represent whether the line is a Spanish accession (purple), MAGIC parent accession (yellow) or MAGIC line (blue). The size of the circles is proportional to the percentage of seeds sown that germinated on a given day. For the two groups of accessions (Spanish accessions and MAGIC parents), examples of the lowest and highest variability lines are shown. For MAGIC lines, examples are shown of low variability (top two lines); high variability, long-tailed (middle two lines) and very high variability bimodal (bottom two lines) lines. **B)** Frequency distribution of coefficient of variation (CV) of germination times for 10 Spanish accessions (purple), the 19 parental natural accessions that were used to generate the MAGIC lines (orange) and 341 MAGIC lines (blue). In the majority of cases, the CV of a given MAGIC line is the mean of the CVs of 3 batches of seeds collected from separate parent plants. **C)** CV of germination times for a subset of 32 MAGIC lines in two separate experiments. The batches of seeds for the two experiments were derived from different independently sown mother plants. The line shows y=x. Figure 1-figure supplement 1 shows the level of reproducibility of germination time distributions across replicates, lengths of period of dry storage, and sowing conditions.

**Figure 1- figure supplement 1.**
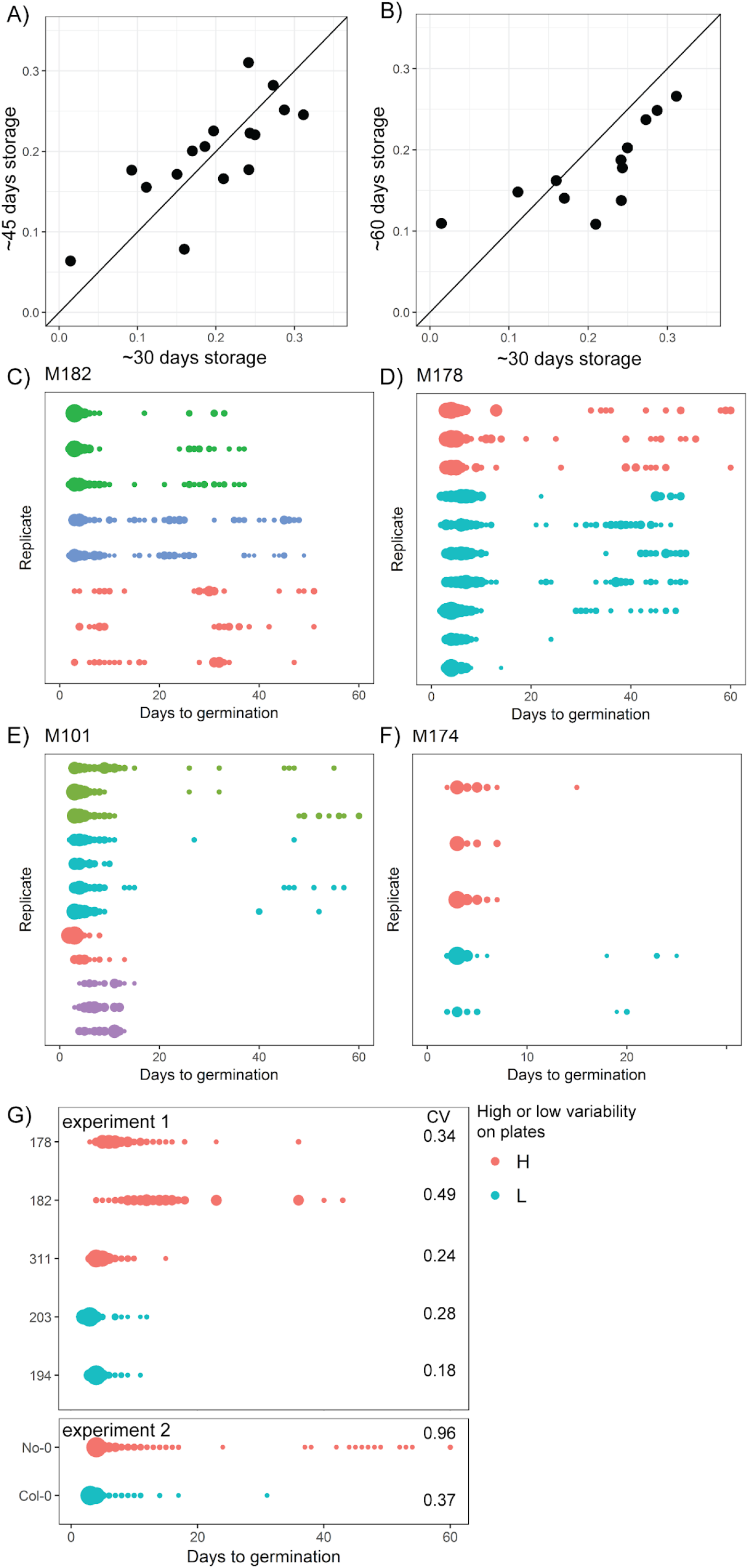
Reproducibility of germination time distributions. **A)** and **B)** CVs of germination time for MAGIC parental accession seeds stored in dry conditions for different lengths of time following harvest (x and y axis labels indicate time of storage). Each point represents a different MAGIC parental accession and is the mean CV across three replicate batches of seeds each collected from a different parent plant. The black lines show x =y. The experimental comparisons are for 16 (A) and 12 (B) of the MAGIC parental accessions. For A, Pearson’s r= 0.77 (95% CI 0.44, 0.91). For B, Pearson’s r= 0.76 (95% CI 0.32, 0.93). **C) - F)** Germination time distributions of the very high variability MAGIC lines shown in Figure 1C. Each row shows the germination time distribution of a seed batch from a different parent plant. Each colour represents seeds collected and sown in a single experiment, with different colours representing different replicate experiments from different parental sowings. The size of the circles is proportional to the percentage of seeds that germinated on a given day. M101 and M174 showed a very late germinating fraction of seeds in some experiments but not others, whilst bimodal distributions were more consistently detected for M182 and M178. **G)** Distributions of germination times on soil for genotypes that were either high or low variability when sown on petri-dishes. Within an experiment, lines that were more variable on plates were also more variable on soil, with the exception of M311.

The accessions showed a range of variabilities (Figure 1A). Some low CV accessions consistently germinated within 4 days, whilst higher CV lines germinated over a period of 19 days (Figure 1A). The MAGIC lines exhibited transgressive segregation, with much greater variation in CV than the parental accessions (Figure 1A, compare orange and blue distributions, Figure 1B). The range of CVs observed in the Spanish accessions was within the range observed across all MAGIC lines (Figure 1B). A small number of MAGIC lines (8 out of 341 characterised) had very high CVs of germination time (>0.6) compared to the rest (Figure 1B), which was due to a fraction of seeds germinating very late, giving rise to bimodal distributions (e.g. M178 and M182 in Figure 1A).

The CV of most MAGIC lines tested was similar between repeat experiments involving independent seed harvests and sowing (Figure 1C) (Pearson’s r = 0.88, 95% CI [0.76, 0.94] for all lines for which repeats were done). In some of the very high variability lines, the presence of very late germinating seeds was reproducible between experiments (e.g. Figure 1C and Figure 1-figure supplement 1, M182 and M178). In other lines, very late germinating seeds were not detected in all experiments, and thus the CV was higher in some experiments than others (e.g. Figure 1C and Figure 1-figure supplement 1, M101 and M174). Thus, although the variability in seed germination time is reproducible for most lines, for some it is possible that their CVs may have been underestimated due to a failure to detect very late germinating seeds. To check whether the level of variability in seed germination time that we obtained for a given line was related to the specific sowing conditions, we sowed selected high and low variability lines on soil and found that although the exact distributions differed slightly between petri-dishes and soil, those lines with higher variability on petri-dishes also had higher variability on soil (Figure 1-figure supplement 1G). Overall our results reveal variation in germination time variability in Arabidopsis, with CVs ranging from 0.09 to 1.7 across the MAGIC lines, giving a good basis for testing the genetic mechanisms underlying this trait.

### Variability in germination times is observed within single siliques

Because germination was characterised for seed samples taken from whole plants, it is possible that the high variability observed in some lines is due to different siliques (fruits) having different germination behaviours. This could arise due to differences in the ages of siliques at the time of seed harvest, or due to positional effects on the parent plant. To address this possibility, we collected seed from individual siliques from 4 high or very high variability lines and characterised their germination time distributions. For these lines, the full range of germination times observed in whole-plant samples was also present in seed from individual siliques (Figure 2A, B; Figure 2-figure supplement 1). This suggests that variability in seed germination time is not an effect of positional or age differences between siliques.

**Figure 2.**
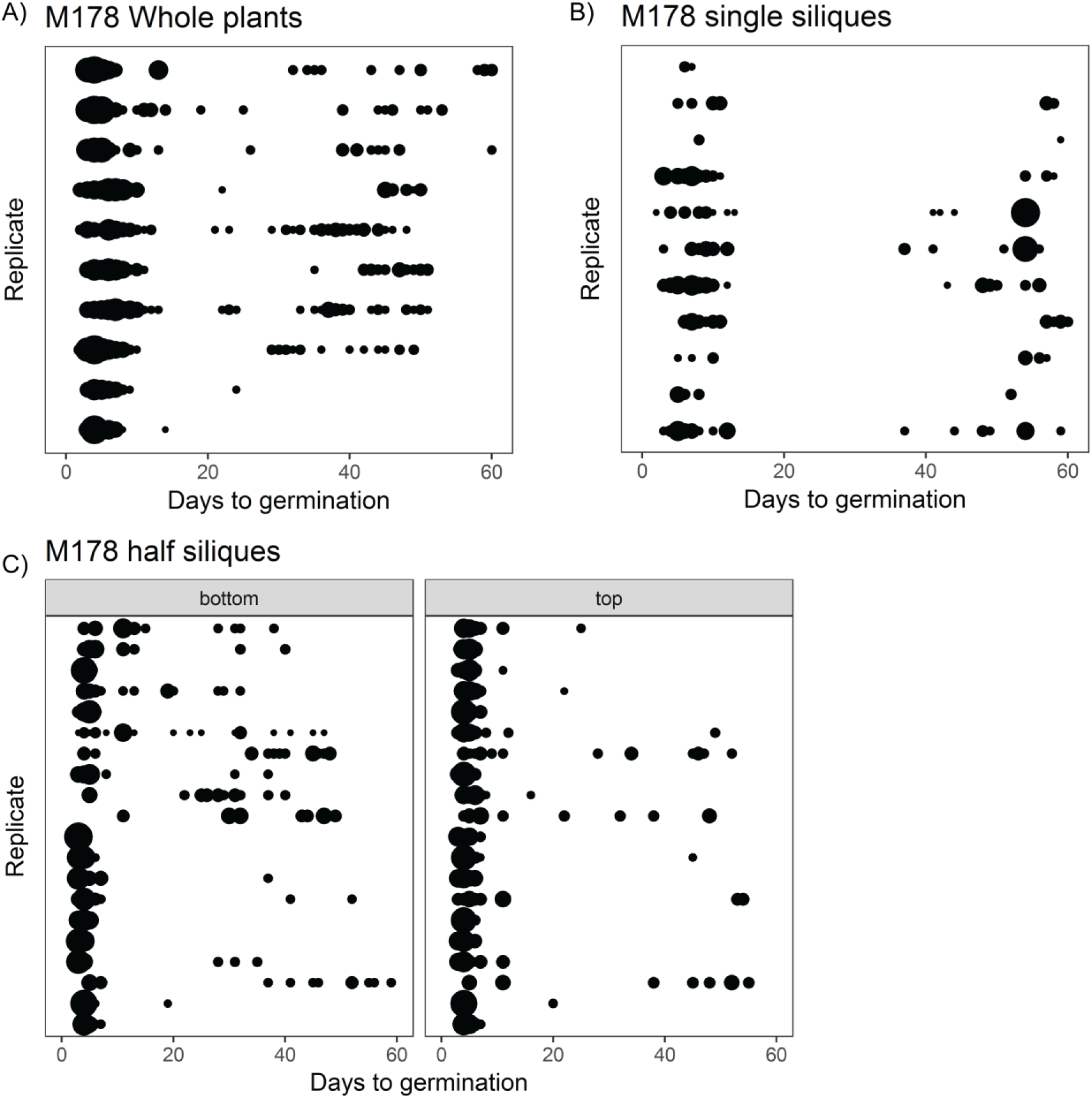
The full range of germination times can be found in individual siliques. **A)** Germination time distributions for a very high variability line, M178. Each row is the distribution obtained using a sample of pooled seeds from one plant, with different rows showing data from different mother plants. **B)** As for A) but each row represents the distribution obtained using seeds from a single silique. **C)** Individual siliques were cut in half and seeds from the top and bottom halves (furthest and closest to the mother plants, respectively) were sown separately. Each row is the bottom and top half of a particular silique. Seeds from whole plants, single siliques and half siliques were obtained and sowed in different experiments. The size of the circles is proportional to the percentage of seeds that were sown that germinated on a given day. Figure 2-figure supplement 1 shows examples for other MAGIC lines plus an experimental repeat and statistical analysis.

**Figure 2-figure supplement 1.**
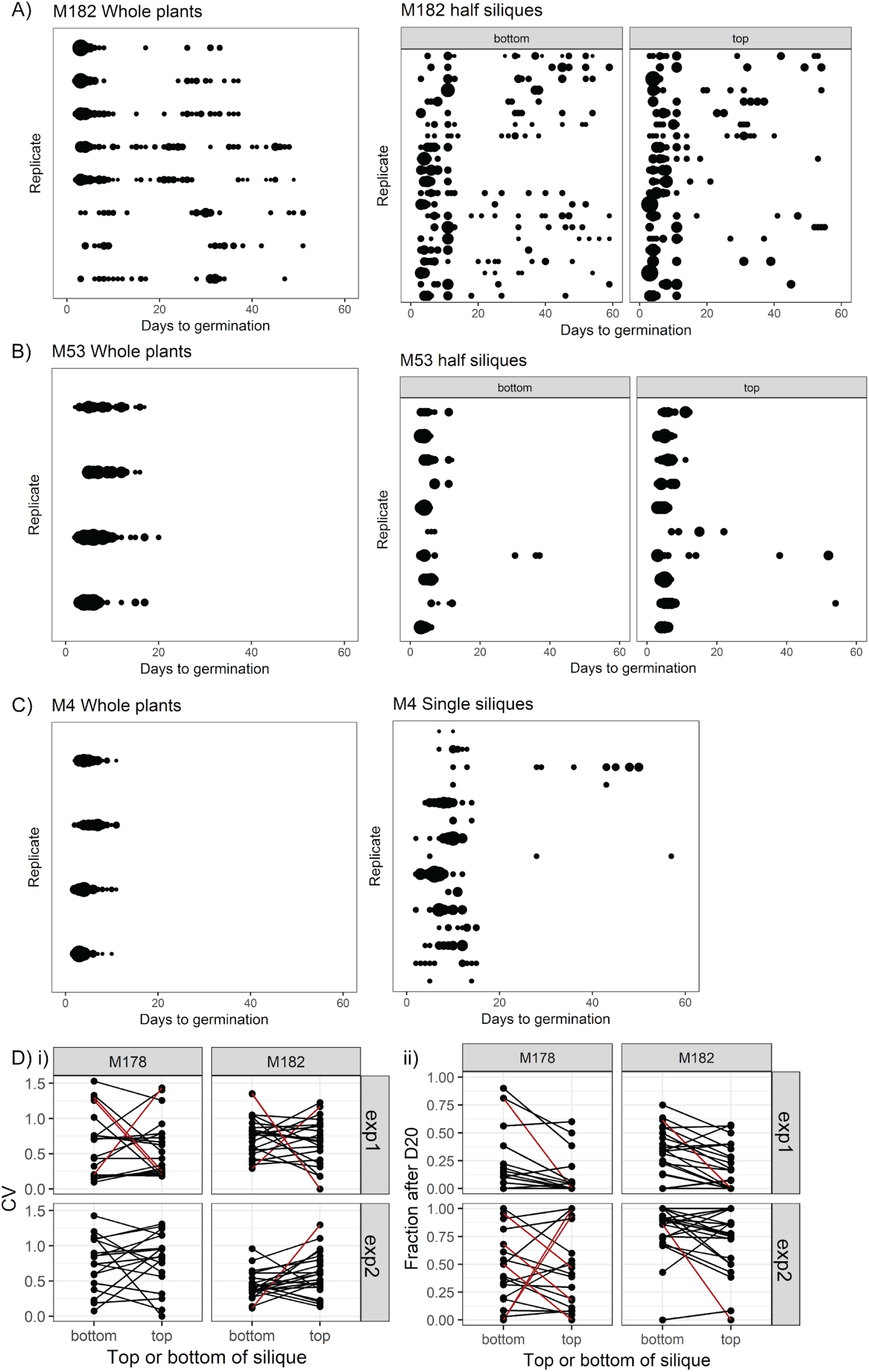
Germination time distributions for whole plants and single siliques for high variability lines. Germination time distributions for M182 **(A)** and M53 **(B)**, for samples of seeds pooled from whole plants, or for single siliques separated into top and bottom halves. Seeds from whole plants and half siliques were obtained and sown in different experiments. For whole plants, each row is the distribution obtained using a sample of pooled seeds from one plant. For half siliques, each row is the bottom and top half of a particular silique. The size of the circles is proportional to the percentage of seeds that were sown that germinated on a given day. **C)** As for A) and B) but for M4, showing whole siliques in the right hand panel. **D)** Comparison of seed germination times between top and bottom halves of individual siliques for M182 and M178 MAGIC lines, collected and sown in two independent experiments (exp1 and exp2). Panels show the CV (panel i) and fraction of late germinating seeds (after day 20, panel ii) from the two halves of each assayed silique. Brown lines indicate significant differences from the null hypothesis (< 5% false-discovery rate). This was determined from bootstrap-based tests that accounted for the unequal number of seeds sown for each silique’s half. Note that even when there is a significant difference between the two silique halves, it is not consistently in the same direction.

We next hypothesised that germination time might be related to the position along the main axis of individual siliques. To test this, we cut siliques into halves and sowed seeds from the top (furthest from the mother plant) and bottom halves separately. For the lines tested, late and early germinating seeds were produced by both halves of the siliques, with no consistent differences between the top and bottom halves in the fraction of seeds that germinated late (Figure 2C; Figure 2-figure supplement 1D). Thus, variability in germination time in the lines tested cannot be explained by positional or maturation gradients within the whole plant or individual siliques. This suggests that a mechanism exists to generate differences in germination behaviour of equivalent seeds from the same silique, which is not dependent on gradients of regulatory molecules along the fruit.

### Variability can be uncoupled from modal germination time and percentage germination

To investigate which types of mechanism might underlie variability in the MAGIC population, we looked at the extent to which variability is correlated with the modal time taken to germinate and the percentage germination within the experiment. For each line, the experiment was defined as complete two weeks after no further seed germinated. If high CV strongly correlates with late germination, or with low percent germination, this would suggest that increased variability in germination times might be simply an effect of differences in average seed dormancy levels between MAGIC lines. If high CV occurs without high time to germination or low percent germination, this would suggest that different lines have different levels of variability independent of their average seed dormancy levels.

We found only a weak correlation between CV and either mode or percent germination, with lower variability lines (low CV) tending to have a lower mode days to germination and higher percentage germination (Figure 3A, Figure 3-figure supplement 1A) (CV *versus* mode, Pearson’s r = 0.27, 95% CI [0.17, 0.37]; CV *versus* percent germination, Pearson’s r = - 0.24, 95% CI [-0.34, −0.14]). Thus, some high variability lines had overall later germination and lower percentage germination than low variability lines, suggesting that they were generally more dormant (Figure 3D, E). However, there were lines that had the same mode days to germination, with very different CVs (Figure 3B, C) and *vice-versa*, lines with the same CV showed a range of modes (Fig 3A ii). There were also lines that were very similar with respect to both percentage germination and mode days to germination, but that had very different CVs (Figure 3-figure supplement 1B, C). Thus, within the MAGIC population, variability can be uncoupled from percentage germination and modal germination time, with low correlation between these traits. The same trends were observed in the natural accessions, where CV was correlated with mode and percentage germination but accessions could be found with similar mode and percentage germination and different CVs (Figure 3-figure supplement 2), raising the possibility that these traits could be under independent selection in nature.

**Figure 3.**
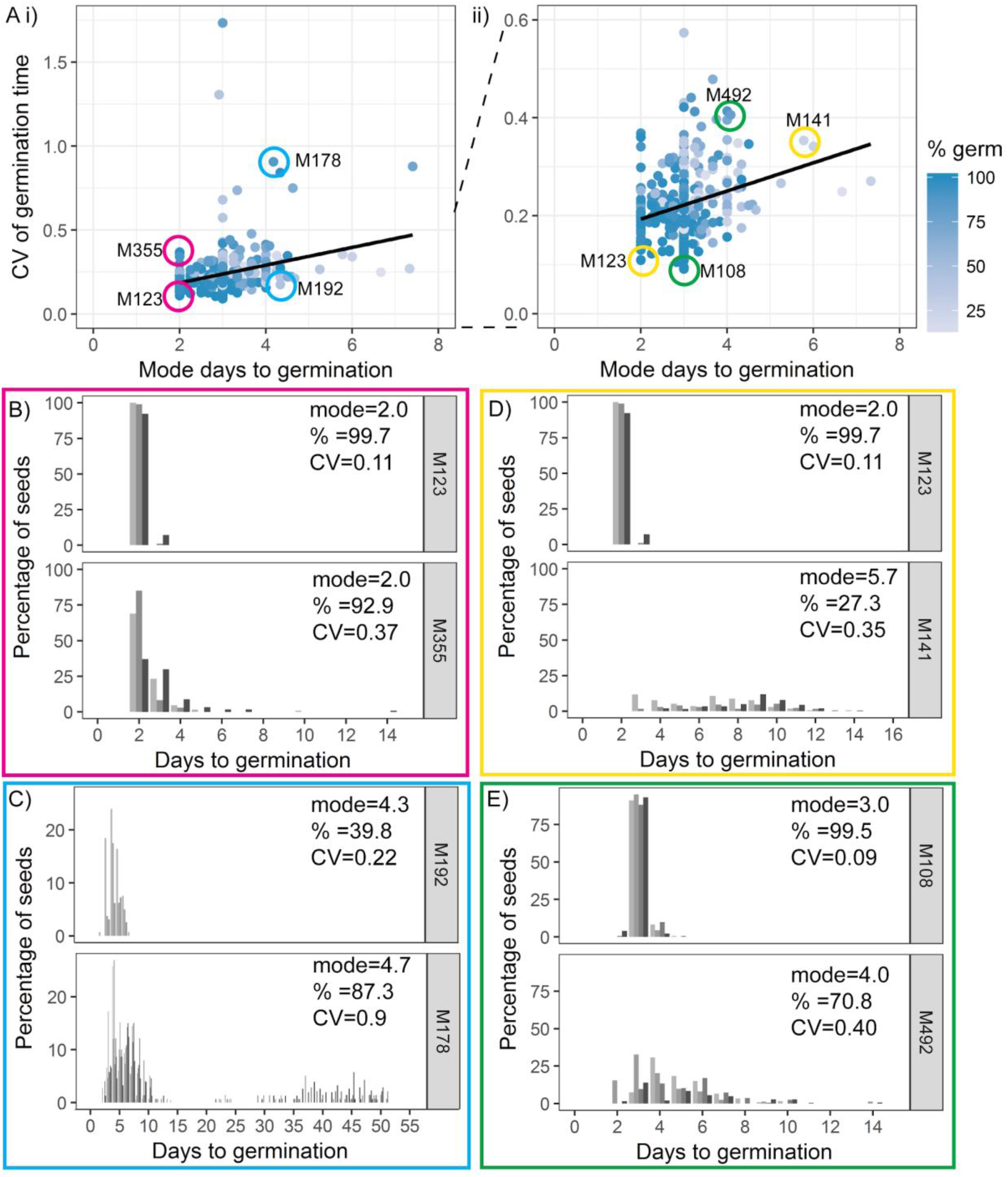
Variability can be uncoupled from modal germination time. **A)** Scatter plots of CV of germination time *versus* mode days to germination for 341 MAGIC lines. Each point is a specific MAGIC line, and in the majority of cases, the CV and mode are mean values obtained from sowing one batch of seeds from each of three separate parent plants. Each point is shaded according to the percentage germination of the line (see scale bar). Coloured circles and labels indicate lines for which examples are shown in B-E. ii) is a zoom in of i) including only lines with CV <0.6. Black lines are linear regressions. **B-E)** Distributions of germination times for pairs of MAGIC lines. The colour of the box matches the coloured circles in A). Lower CV lines are shown on top. Grey coloured bars show the germination distribution of seed batches from replicate mother plants. **B** and **C)** Exemplar lines with the same mode days to germination but different CVs of germination time. **D** and **E)** Lines that have different CVs and different mode days to germination. For each line, the mode days to germination, final percentage germination and CV of germination time are shown. Note that the x-axis scale differs between plots. Figure 3-figure supplement 1 shows the relationship between CV and percentage germination for MAGIC lines. Figure 3-figure supplement 2 shows relationships between CV, mode and percentage germination for natural accessions.

**Figure 3-figure supplement 1.**
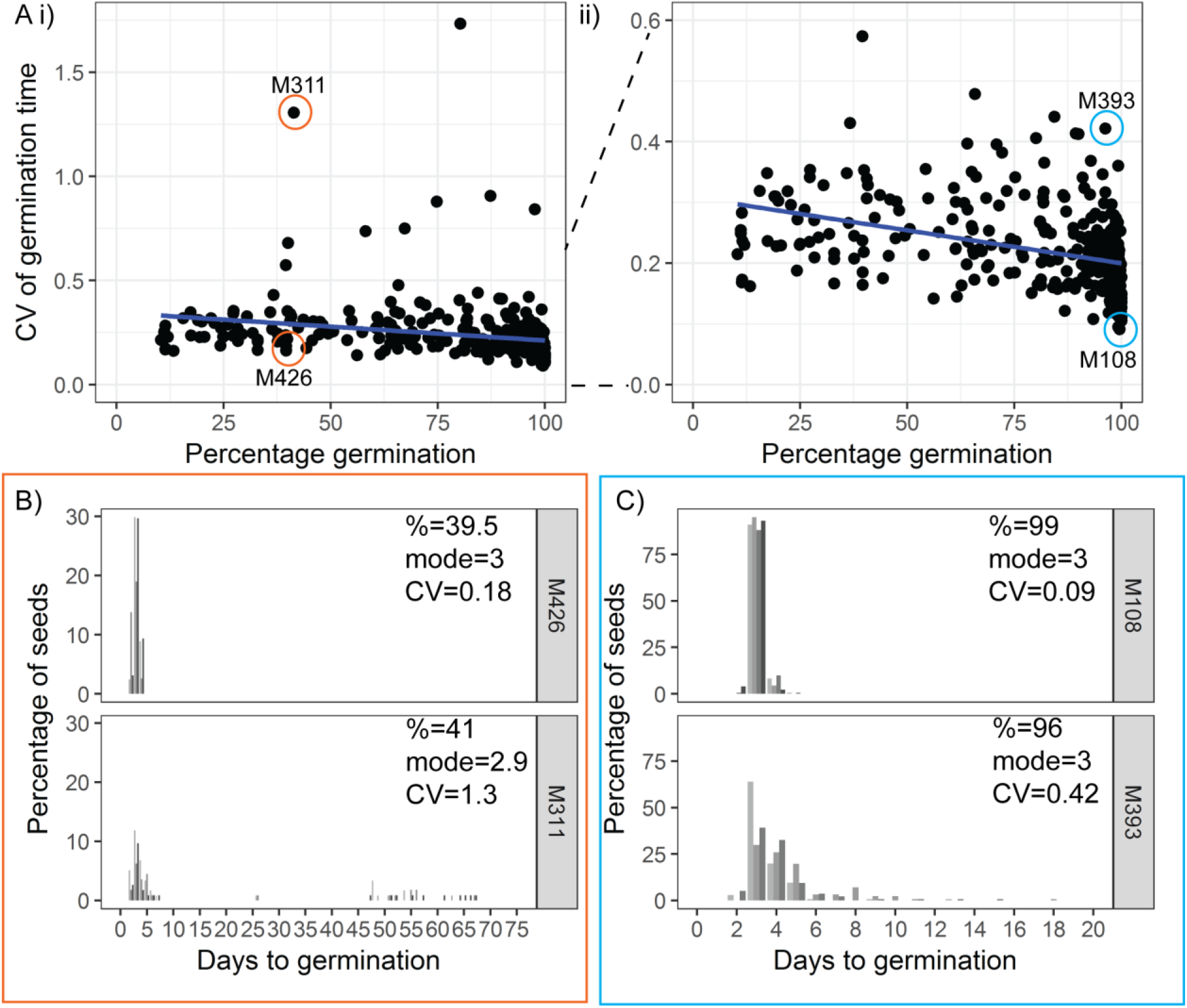
Variability can be uncoupled from percentage germination. **A)** Scatter plots of CV of germination time, *versus* percentage germination, for 341 MAGIC lines. Each point is a specific MAGIC line, and in the majority of cases, the CV and percent are mean values for three batches of seeds from separate parent plants. Coloured circles and labels indicate lines for which examples are shown in B-C. ii) is a zoom in of i), including only lines with CV <0.6. Blue lines are linear regressions. **B-C)** Distributions of germination times for exemplar MAGIC lines that have similar final percentage germination and mode days to germination but different CVs. The colour of the box matches the lines to the coloured circles in A. Lower CV lines are shown on top. Grey coloured bars show the distribution of germination time for seed batches from replicate mother plants. Note that the x-axis scale differs between B and C.

**Figure 3-figure supplement 2.**
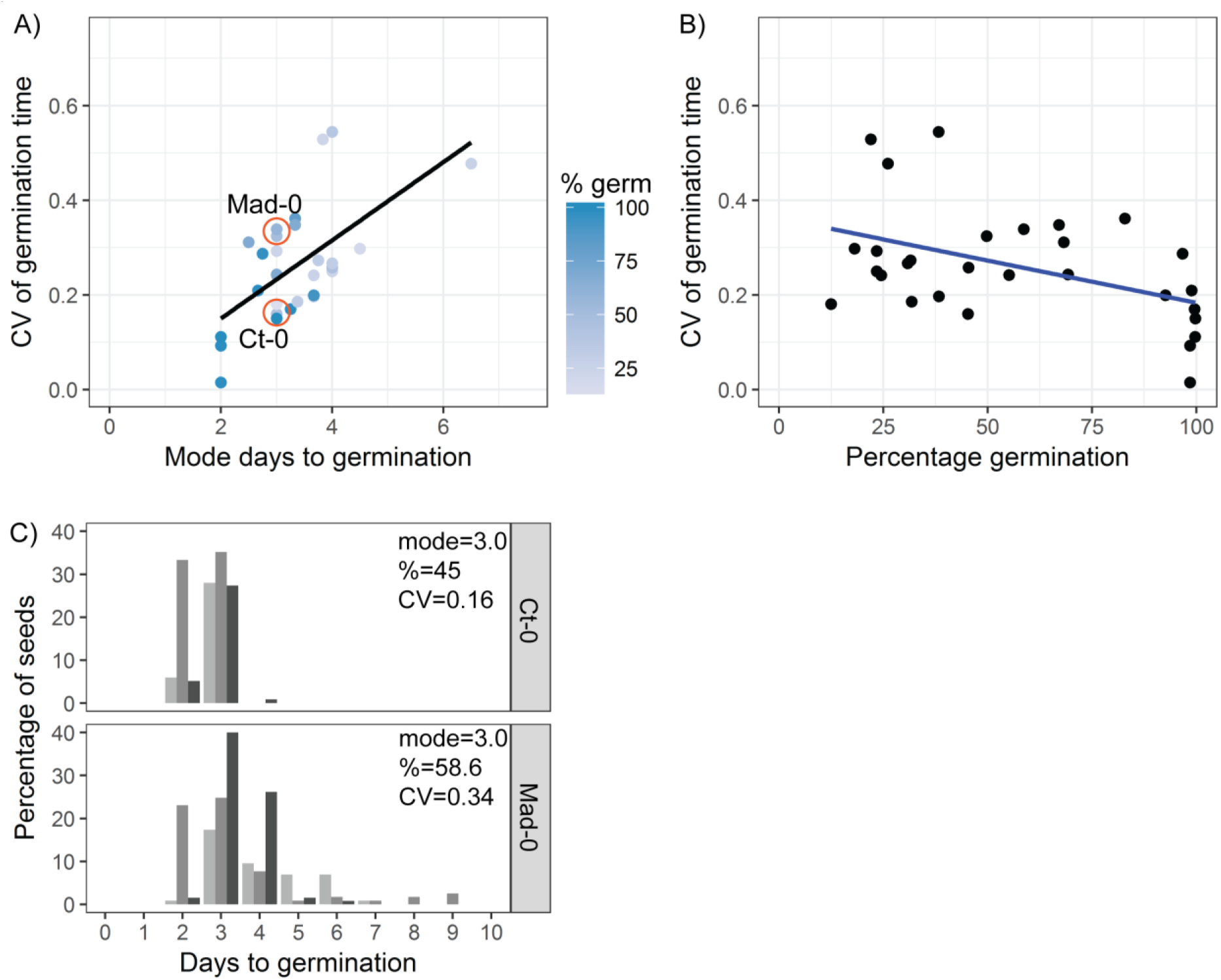
The relationship between CV, mode days to germination and percentage germination in natural accessions. **A)** Scatter plot of CV of germination time *versus* mode days to germination for the 19 parental accessions of the MAGIC lines and 10 Spanish accessions. Each point is a specific accession, and the CV and mode are means for one batch of seed from each of at least three separate parent plants. Each point is shaded according to the percentage germination of the line (see scale bar). Orange circles indicate lines for which examples are shown in C. Trend line is a linear regression. For CV *versus* mode, Pearson’s r = 0.61, 95% CI [0.31, 0.79]. **B)** As for A, but showing CV *versus* percentage germination. For CV *versus* percent germination, Pearson’s r = - 0.46, 95% CI [-0.71, −0.11]. **C)** Distribution of germination times for exemplar accessions, with similar mode and percentage germination but different CVs (accessions indicated in A). Grey coloured bars show the distributions of germination for seed batches from replicate mother plants.

### QTL mapping in MAGIC lines reveals a candidate variability-specific locus

We next performed QTL mapping on the germination data for the MAGIC lines (Kover et al., 2009) to investigate the genetics of germination time variability. The full set of MAGIC lines phenotyped includes lines with different types of germination time distributions. All low variability lines and most high variability lines have unimodal distributions of germination time (e.g. Figure 1A, M108, M203, M305, M393, M285). However, there are 8 lines that tend to have bimodal distributions when sown on agar (e.g. Figure 1A, M182 and M178). As such, these lines lie at the extreme tail of the distribution of CVs, with much higher values than the other lines (Figure 1B). Therefore, we ran our QTL scans both with and without the bimodal lines, as their extreme values may affect the QTL results disproportionately (Figure 4A, B).

**Figure 4.**
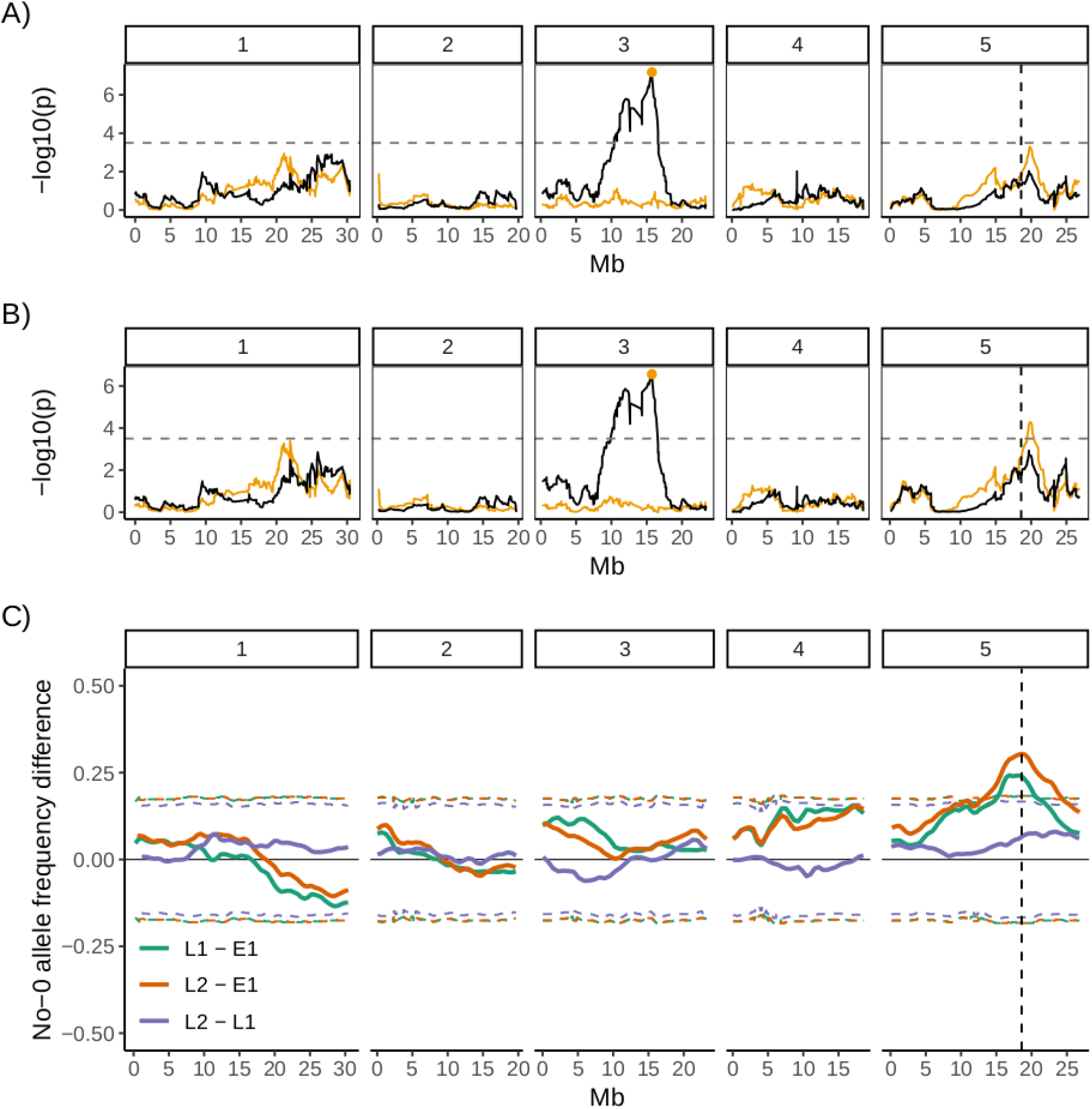
QTL and bulk segregant mapping reveals two QTL underlying CV of germination time. **A)** and **B)** Manhattan plots showing the QTL association results for each marker individually (black line) and for each marker together with the Chr 3 QTL marker added as a covariate in the model (orange line). The marker used as a covariate is highlighted with an orange point. A) is for the full set of 341 MAGIC lines that was phenotyped and B) excludes the 8 bimodal lines with very high CV. The y-axis shows the p-values for the 1254 markers used, on a negative log_10_ scale. The numbered panels represent the 5 chromosomes of Arabidopsis. The horizontal dashed line shows a 5% genome-wide threshold corrected for multiple testing (based on simulations in Kover et al, 2009). The vertical dashed line indicates the *DOG1* gene. Figure 4-figure supplement 1 shows QTL mapping for mean and mode days to germination and percentage germination. Figure 4-figure supplement 2 shows estimated effects of accession haplotypes on CV, mode and percent germination. **C)** Mapping QTL by bulk-segregant analysis using whole-genome pooled sequencing of F2 pools from a Col-0 x No-0 cross. One early and two late germinating F2 pools were sequenced. The plot shows the No-0 allele frequency differences between pairs of pools indicated in the legend (Figure 4-figure supplement 3 shows details of pool selections, E1= early pool, L1= late 1 pool, L2= late 2 pool). The horizontal dashed lines indicate the 95% thresholds based on simulating the null hypothesis of random allele segregation, taking into account the size of the sampled pools and the sequencing depth at each site (Magwene et al., 2011; Takagi et al., 2013). Positive values above the top line indicate enrichment for No-0 alleles, while negative values below the bottom line indicate enrichment for Col-0 alleles. As predicted, late germinating pools were enriched for the No-0 haplotype in the region of the Chr5 QTL. Here the peak of association overlaps with the *DOG1* gene (dashed vertical line). Figure 4-figure supplement 4 shows germination phenotypes of F3 seeds from Col-0 x No-0 F2 plants that themselves germinated early or late.

**Figure 4-figure supplement 1.**
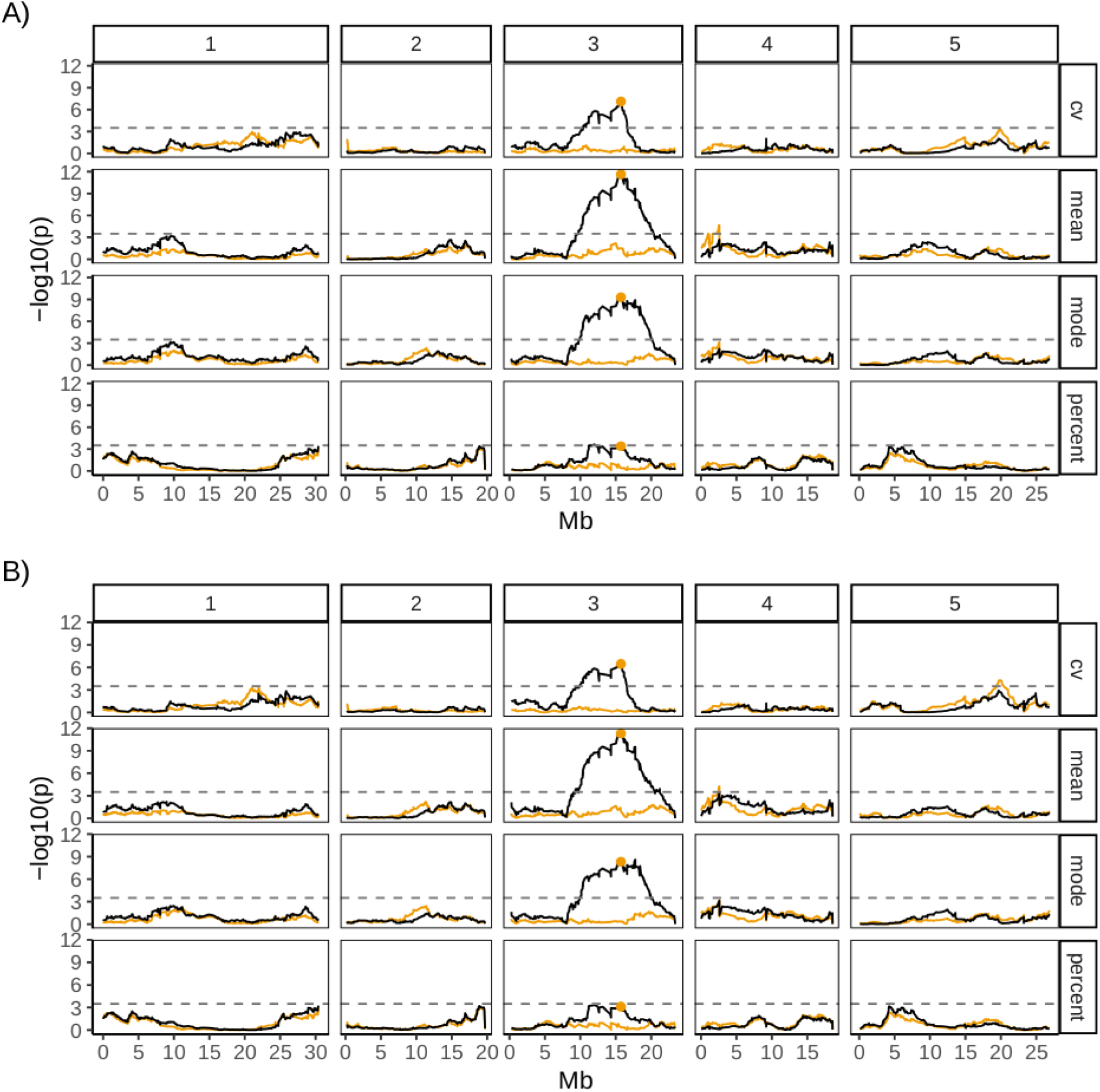
QTL mapping for germination traits, with and without bimodal MAGIC lines. **A)** is for all 341 MAGIC lines that were phenotyped, **B)** is for 333 of these lines (the full set minus the 8 bimodal lines with very high CV). The y-axis shows the p-values for the 1254 markers used, on a negative log10 scale. The numbered panels represent the 5 chromosomes of Arabidopsis and the scan was performed for 4 germination traits: CV of germination time, mean germination time, mode germination time and percentage germination. The horizontal dashed line shows a 5% genome-wide threshold corrected for multiple testing (based on simulations in Kover *et al*., 2009).

**Figure 4-figure supplement 2.**
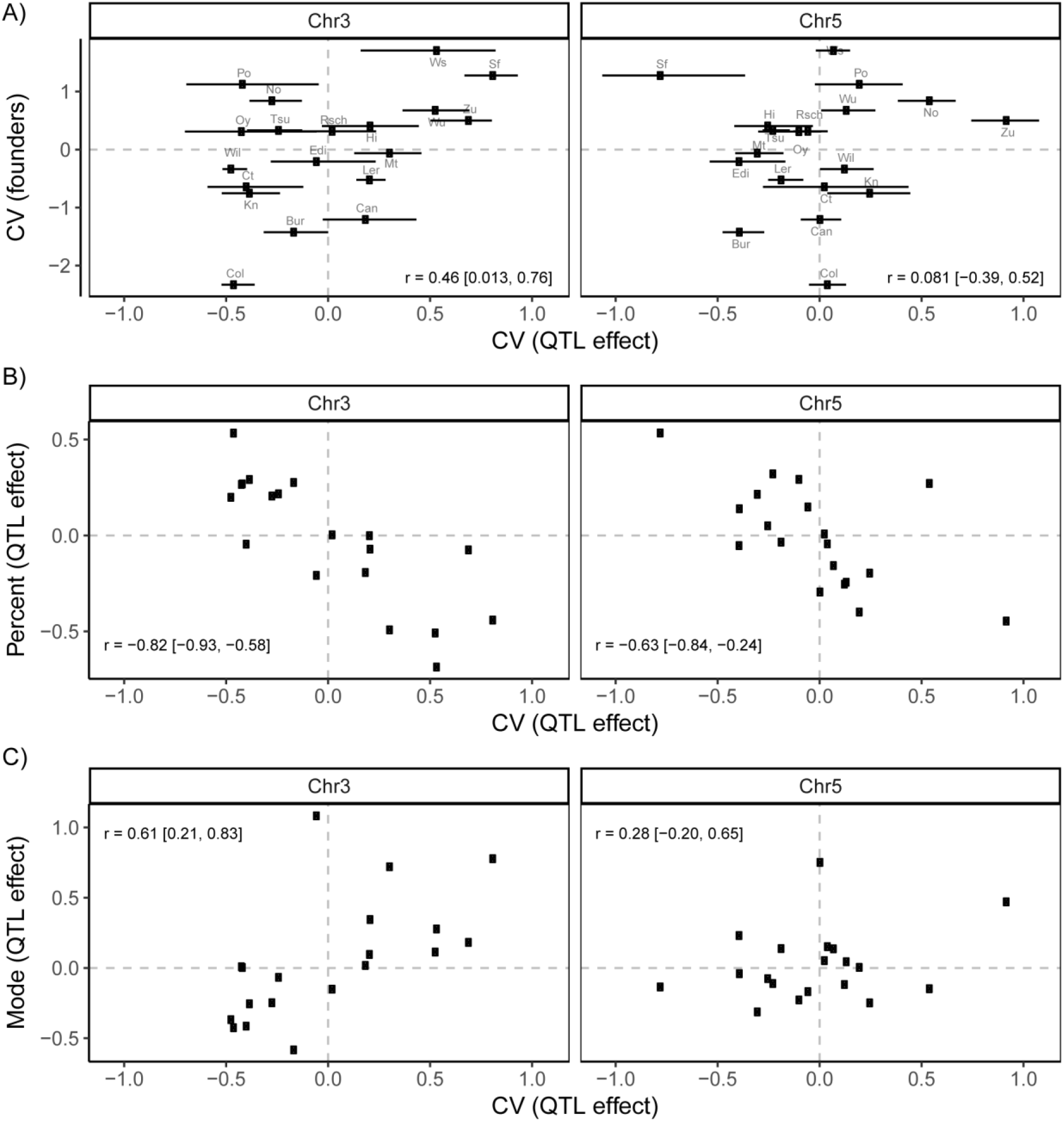
Accession-specific QTL effects on CV, mode and percent germination. **A)** Correlation between the germination CV of the founder accessions and the predicted accession effects at the two putative QTL on Chr 3 and Chr 5 (Fig 4). The effects of the 19 parental accession haplotypes were estimated by calculating the mean CV of MAGIC lines inferred to carry each particular haplotype. **B)** Correlation between predicted QTL effects on percent germination and CV. **C)** Correlation between predicted QTL effects on mode days to germination and CV. In all panels, the mean effect of each parental accession’s QTL allele was estimated from the probabilistic assignment of each MAGIC line to that founder parent (Kover et al 2009). Error bars in panel A) show the 95% confidence intervals of these estimates (these were omitted from the other panels for clarity). All trait values were standardized, so that axis units represent the number of standard deviations away from the respective mean, with the horizontal and vertical dashed lines at zero highlighting the mean of the respective trait in the population. Pearson’s correlation, r, is indicated in each panel, with the 95% confidence interval in brackets.

**Figure 4-figure supplement 3.**
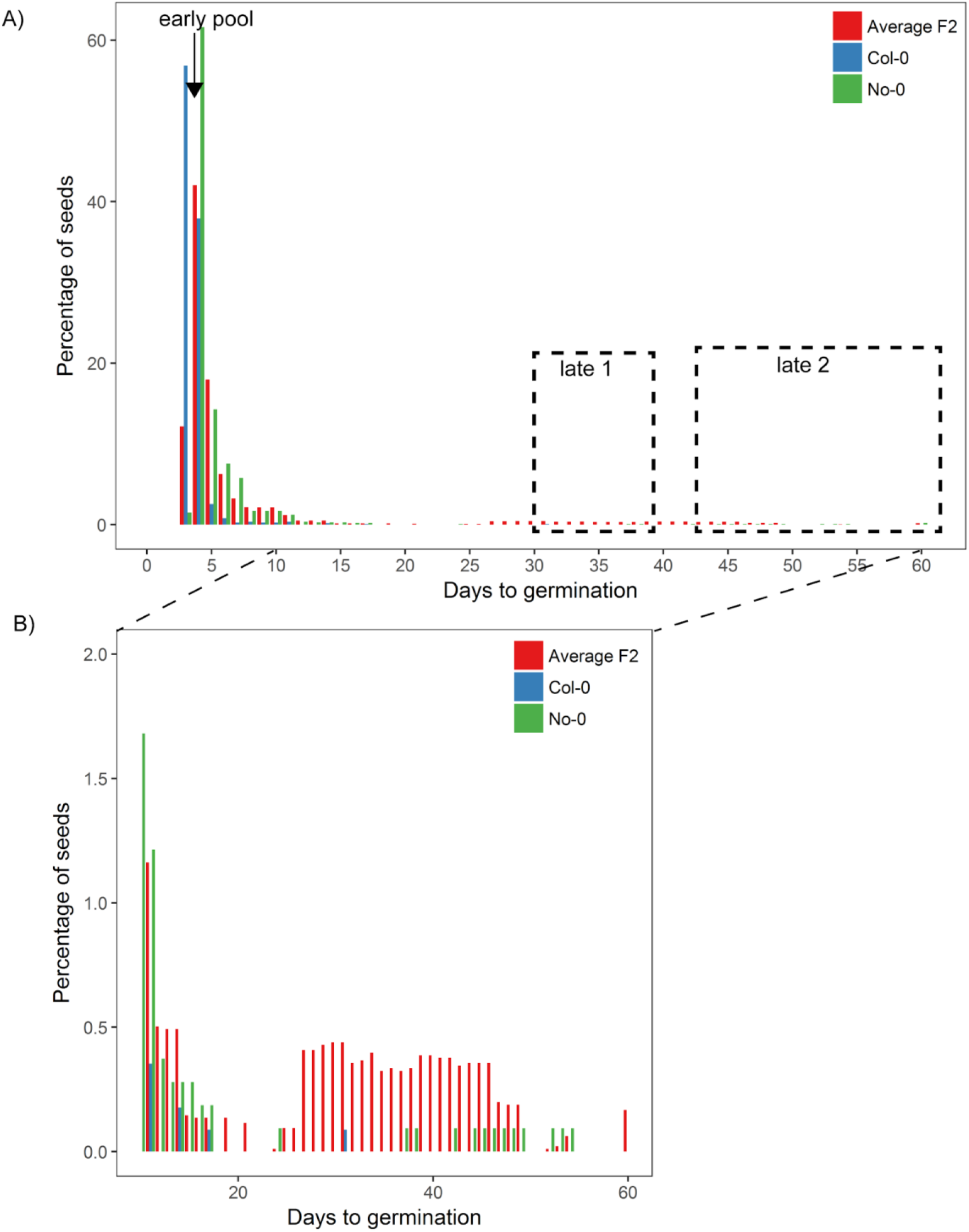
Germination time distributions and DNA-seq pools of Col-0 x No-0 F2. 8 batches of Col-0 x No-0 F2 seeds, each containing ∼1100 seeds and collected from a different F1 parent plant, were sown on soil. Seeds from parental accessions Col-0 and No-0 were also included in the experiment and for these, batches of ∼1100 seeds pooled from 3 parent plants were sown for each accession. The different F2 batches behaved similarly, so here we present an averaged distribution based on bulked data. The percentage germination each day is a percentage of all seeds that germinated (rather than of all seeds that were sown). **A)** shows the full germination time distribution with pools used for DNA sequencing highlighted and **B)** shows days 10-60, with a different y axis scale to show the late germinating seeds. The early pool used for sequencing was composed of 152 individuals that germinated on day 4; the “late 1” pool was composed of 321 individuals that germinated between days 31 and 39 and the “late 2” pool was composed of 213 individuals that germinated between days 43 and 60. We reasoned that, since late germination is predominantly restricted to the more variable parent (No-0), late germinating F2 plants should be enriched for the No-0 accession at loci promoting high variability (including the chr5 locus at ∼20Mb where the No-0 haplotype is predicted to promote high CV).

**Figure 4-figure supplement 4.**
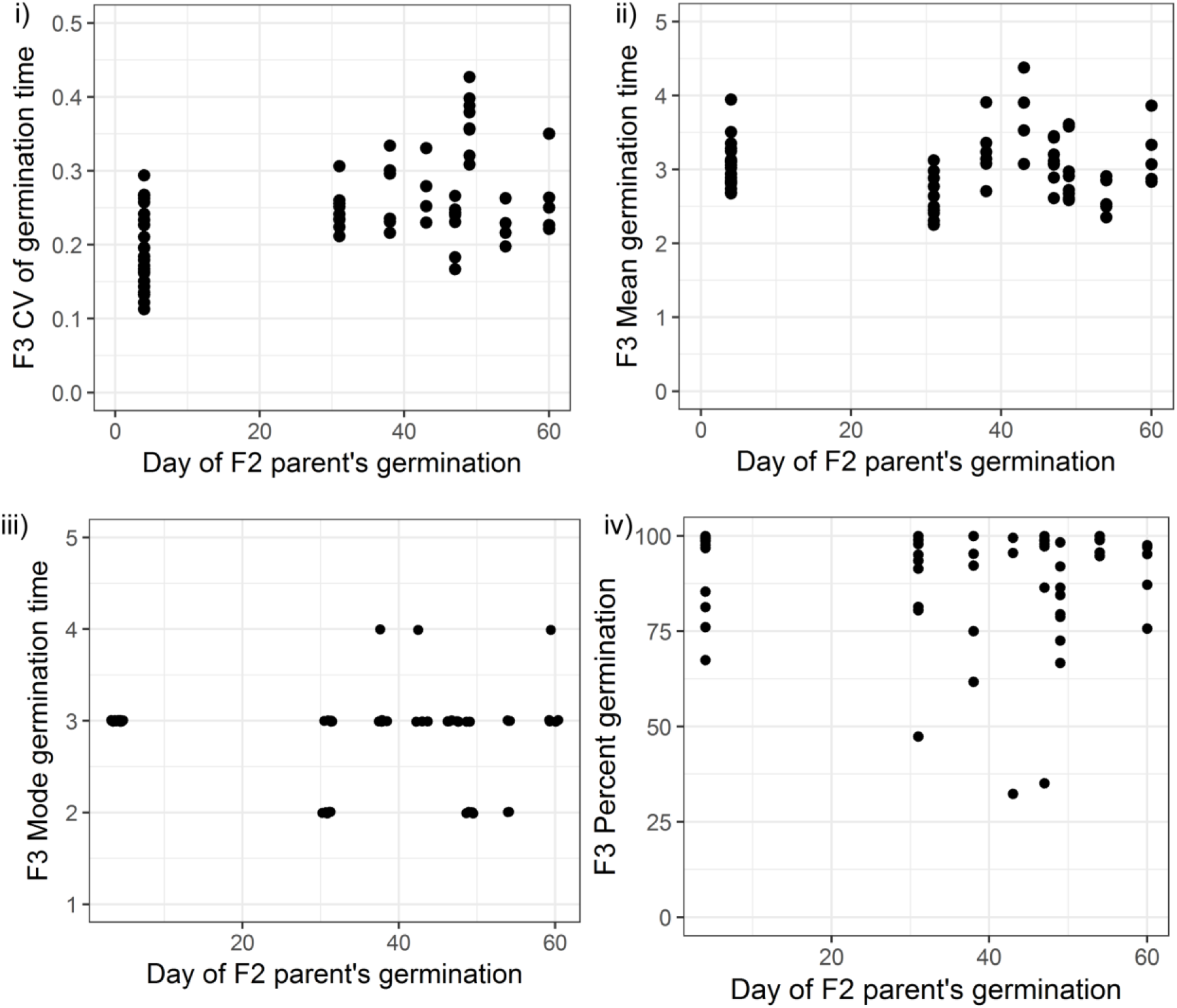
Germination phenotypes of F3 seeds from Col-0 x No-0 F2 parent plants that themselves germinated early or late. CV, mean, mode and percent germination for F3 seeds collected from F2 plants that themselves germinated at different times (x axis). Batches of seeds from F2 plants that germinated late (between days 30 and 60) had, on average, a significantly higher CV than seeds from plants that germinated early (on day 4) (the mean CVs of the two groups were 0.19 in the early group *versus* 0.27 in the late group, Wilcoxon rank sum test W = 199, p-value =1.163e-05, n=23 for seeds from early germinators, n=47 for seeds from late germinators). Seeds of late germinating plants did not tend to have higher mean or mode germination times (the mean of mean germination times was 3.05 days in the early group *versus* 3 days in the late group; the mean of mode germination times was 3.00 in the early group *versus* 2.78 in the late group). Percentage germination shows a small but significant difference between the two groups of F3 seeds (mean percentage germination: 95.3 in the early group *versus* 87.67 in late group, Wilcoxon rank sum test W = 754, p-value = 0.00419, n=23 for seeds from early germinators, n=47 for seeds from late germinators).

QTL mapping for both the full set of lines, and the set excluding outliers, revealed a region of chromosome 3 that accounted for ∼14% of the variance in CV of germination time for the MAGIC lines used (Figure 4A, B). The region of significant association was broad and spanned the centromere. The tip of the peak co-located with the previously identified *DELAY OF GERMINATION 6* QTL (*DOG6*), at 15.9 Mb (Bentsink et al., 2010; Hanzi, 2014). This chromosome 3 QTL was also associated with mean days to germination, mode days to germination and percentage germination, suggesting this locus is a general regulator of germination time, rather than specifically affecting variability (Figure 4-figure supplement 1).

To investigate whether other loci explained any residual variance not explained by this major locus, we ran the QTL scans using the chromosome 3 QTL genotype as a covariate in the model. This revealed a further putative QTL at 19.8 Mb on chromosome 5 associated with CV (Fig 4 B). Unlike the chromosome 3 locus, this one showed no association with mode or mean days to germination and was not significantly associated with percentage germination (Figure 4-figure supplement 1), suggesting that it may specifically regulate variability independently from the average germination time and level of dormancy. This locus accounts for an extra 9% of the variance in CV of the MAGIC lines used. The QTL peak lies ∼1.2 Mb downstream of the *DELAY OF GERMINATION 1* (*DOG1,* AT5G45830) gene (at 18.59 Mb) and ∼1.2 Mb upstream from the *Seedling Emergence Time 1 (SET1)* locus (at ∼21 Mb) (Footitt et al., 2019). The QTL scans with and without the bimodal lines were very similar for the four germination traits (CV, mean, mode days to germination and percentage germination) except the Chr5 peak was not significantly associated with CV when bimodal lines were included (Fig 4A; Figure 4-figure supplement 1).

We next estimated the effects of particular accession haplotypes at the two QTL on the different germination traits (Figure 4-figure supplement 2). This revealed that for the Chr3 QTL, but not for the Chr 5 QTL, there was a weak positive correlation between the CV of germination time of the parental accessions and the estimated effect of their QTL haplotypes on CV in the MAGIC lines (Figure 4-figure supplement 2A). For the chromosome 3 QTL, there was a relatively strong negative correlation between haplotypic effects on CV and percent germination, and a positive correlation between effects on CV and mode (Figure 4-figure supplement 2B-C). This supports the conclusion that this QTL is a general regulator of seed germination time. For the chromosome 5 QTL, there was a negative correlation between haplotypic effects on CV and percent germination, but there was no correlation between their effects on CV and on mode (Figure 4-figure supplement 2B-C). This suggests that the chromosome 5 QTL may influence variability independently of the average time to germination, but may also have an effect on the percentage germination.

To confirm the effect of the chromosome 5 QTL on CV in an independent experiment, we used an F2 bulked segregant mapping approach in a cross between two accessions (Col-0 and No-0) predicted to have haplotypes in this genomic region with different effects on CV (Figure 4-figure supplement 2 A, Chr5 panel). We performed whole genome sequencing on pools of F2 plants that germinated late, and so were predicted to be enriched for the No-0 haplotype at ∼20Mb on Chr5, promoting high CV, and compared their sequences to those of a pool of early germinating F2 plants (Figure 4C, for details of pools see Figure 4-figure supplement 3). The results independently verified that a locus at ∼20Mb of Chr5 has an influence on CV. In this experiment, the peak of association was located at 18.6Mb on Chr5, which overlaps precisely with the *DOG1* gene (Figure 4C). We also quantified germination traits of the F3 offspring of F2 plants that themselves germinated early or late. This showed that late germinating F2 plants produced seed with higher CVs of germination time, lower percentages of germination and similar average germination times compared to seeds of early germinating F2 plants (Figure 4-figure supplement 4).

In summary, we have shown that at least two loci contribute to variability in seed germination time in the MAGIC lines (chromosome 3, ∼16 Mb and chromosome 5, ∼18.6/ 19.8 Mb). Although variability can be separated from mode days to germination and percentage germination, the main QTL on chromosome 3 has correlated effects on all these three traits. The locus at ∼19 Mb on chromosome 5 appears to affect variability independently of average time to germination and likely accounts for some of the variation in CV that occurs even for lines with the same mode days to germination.

The chromosome 5 peak obtained in the bulk segregant mapping overlaps with the *DOG1* gene known to play a role in seed dormancy level. The peak obtained in the QTL mapping is slightly shifted and lies equidistant between *DOG1* and the nearby *SET1* locus (at ∼21Mb) which affects dormancy levels in the field in response to environmental conditions. Consistent with a role for this region of chromosome 5 in seed dormancy in the MAGIC lines, its haplotypic effects on CV and on percentage germination were negatively correlated (Figure 4-figure supplement 2 B, Chr5 panel). Additionally, our Col-0 x No-0 F2 and F3 analysis suggested that seeds from plants enriched for the No-0 haplotype at this locus (which is associated with high CV) had a lower percentage germination than seeds from plants enriched for the low CV Col-0 haplotype (Figure 4-figure supplement 4). However, perhaps surprisingly, this locus was not significantly associated with percentage germination in the QTL mapping (Figure 4A, B). This may be because, unlike the Cvi accession that was used originally to map both *DOG1* and *SET1* loci (Alonso-Blanco et al., 2003; Footitt et al., 2019) the accessions used to generate the MAGIC lines have relatively weak dormancy and may not carry alleles in this region that promote dormancy sufficiently strongly to be detected in the QTL mapping.

### A stochastic model of the GA/ABA bistable switch can account for the observed genetic variation in germination time distributions

There is evidence to suggest that candidate genes underlying our identified loci are related to ABA signalling or sensitivity. The effect of the *DOG6* locus overlapping our chromosome 3 QTL is thought to be caused by the *ANAC060* gene, which regulates ABA sensitivity (Hanzi, 2014; Li et al., 2014). Both the *DOG1* gene and the *SET1* locus possibly underlying the chromosome 5 peak have been proposed to modulate ABA signalling (Footitt et al., 2019; Fuchs et al., 2013; Née et al., 2017). This raises the question of how genetic loci could have different effects on germination time distributions by influencing the ABA pathway. To answer this, we built a simplified mathematical model of the core ABA-GA network that governs germination time (Liu and Hou, 2018) (Figure 5A). We wished to understand whether different parts of the network could be modulated to have either correlated or distinct effects on CV, mode and percentage germination.

**Figure 5.**
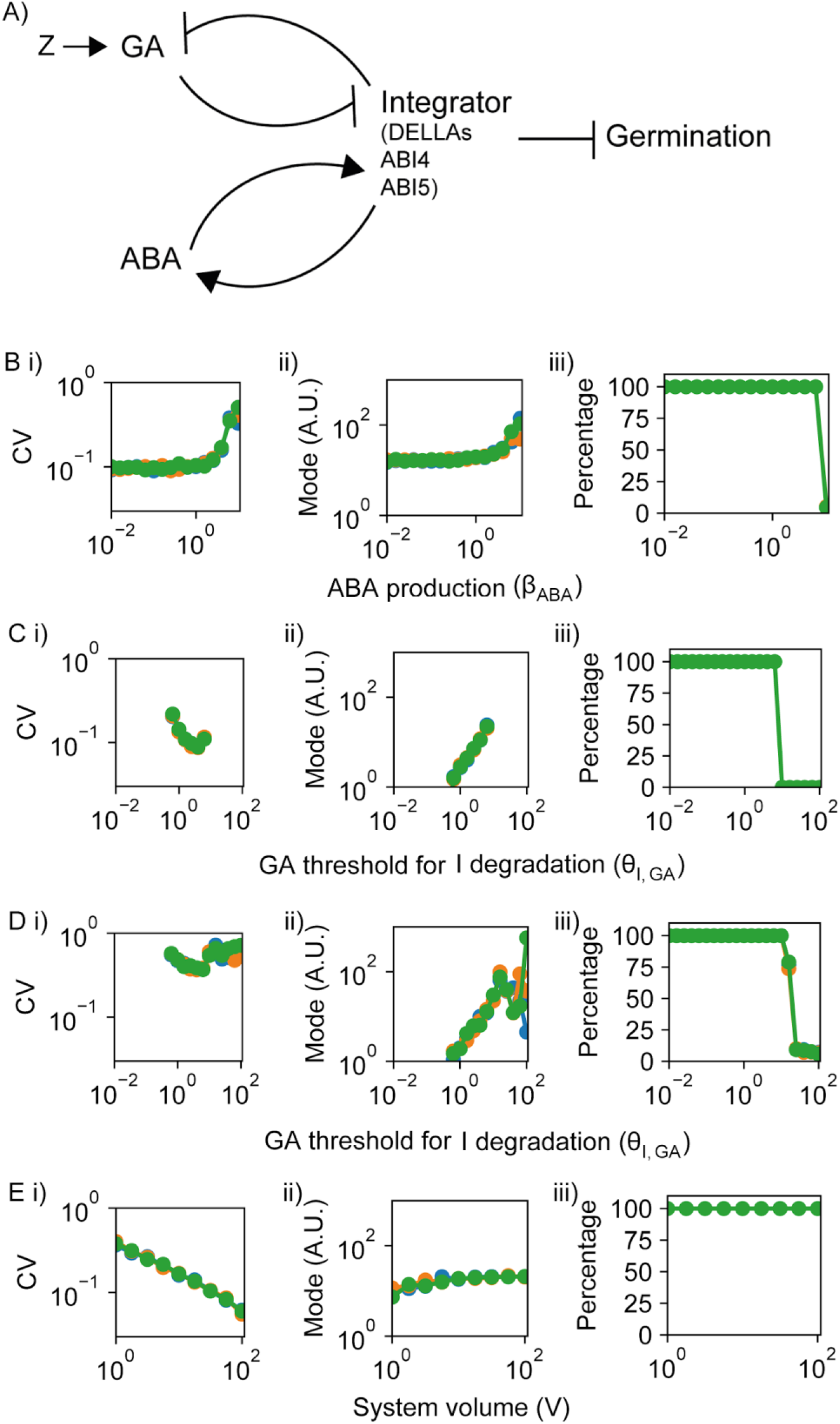
Model of the ABA-GA bistable switch and effects of its parameters on germination traits. **A)** Model of the ABA-GA network. We represent the inhibitors of germination - DELLAs, ABI4 and ABI5 - as one factor, called Integrator, which we assume must drop below a threshold for germination to occur. We assume that ABA promotes the production of Integrator, and that GA promotes its degradation. Integrator is assumed to promote ABA production and inhibit GA production. A factor, Z, increases upon sowing and promotes GA biosynthesis. **B)-E)** show the effects on CV, mode and percentage germination of simulated germination time distributions as single parameter values are changed. Each panel shows the results of three different runs of stochastic simulations on 400 seeds, represented in different colours. **B)** Varying the rate of ABA production as an example of a parameter that, when changed, tends to have positively correlated effects on CV and mode of germination time. **C)** Varying the threshold of GA for degradation of Integrator (this parameter is inversely correlated with sensitivity of Integrator to GA). For some points in parameter space, varying this parameter has anti-correlated effects on CV and mode. **D)** As for C, but in a different region in parameter space (see below for parameter details), in which increasing the threshold of GA for degradation of Integrator causes an increase in mode with relatively constant CV. **E)** Varying the effective system volume parameter, V, which controls the level of noise in the system (noise intensity is proportional to 1/V), as an example of a parameter that, when changed, causes decoupled effects on CV and mode. For some areas of parameter space an increase in V, and therefore a decrease in noise, causes the CV to decrease but leaves mode and percentage germination relatively unchanged. Parameter values are provided in the methods and were the same across simulations with the exception of the parameters varied on x axes and differences in *v_ABA_* in B), θ*_I,ABA_* in C) and E) and V in D). Figure 5-figure supplement 1 provides information on the dynamics of the model. Figure 5-figure supplement 2 shows simulated germination time distributions corresponding to the parameter explorations in (B-E). Figure 5-figure supplement 3 shows the full results of the 2D parameter screen in terms of the effects of parameter pairs on CV, mode and percentage germination.

**Figure 5-figure supplement 1.**
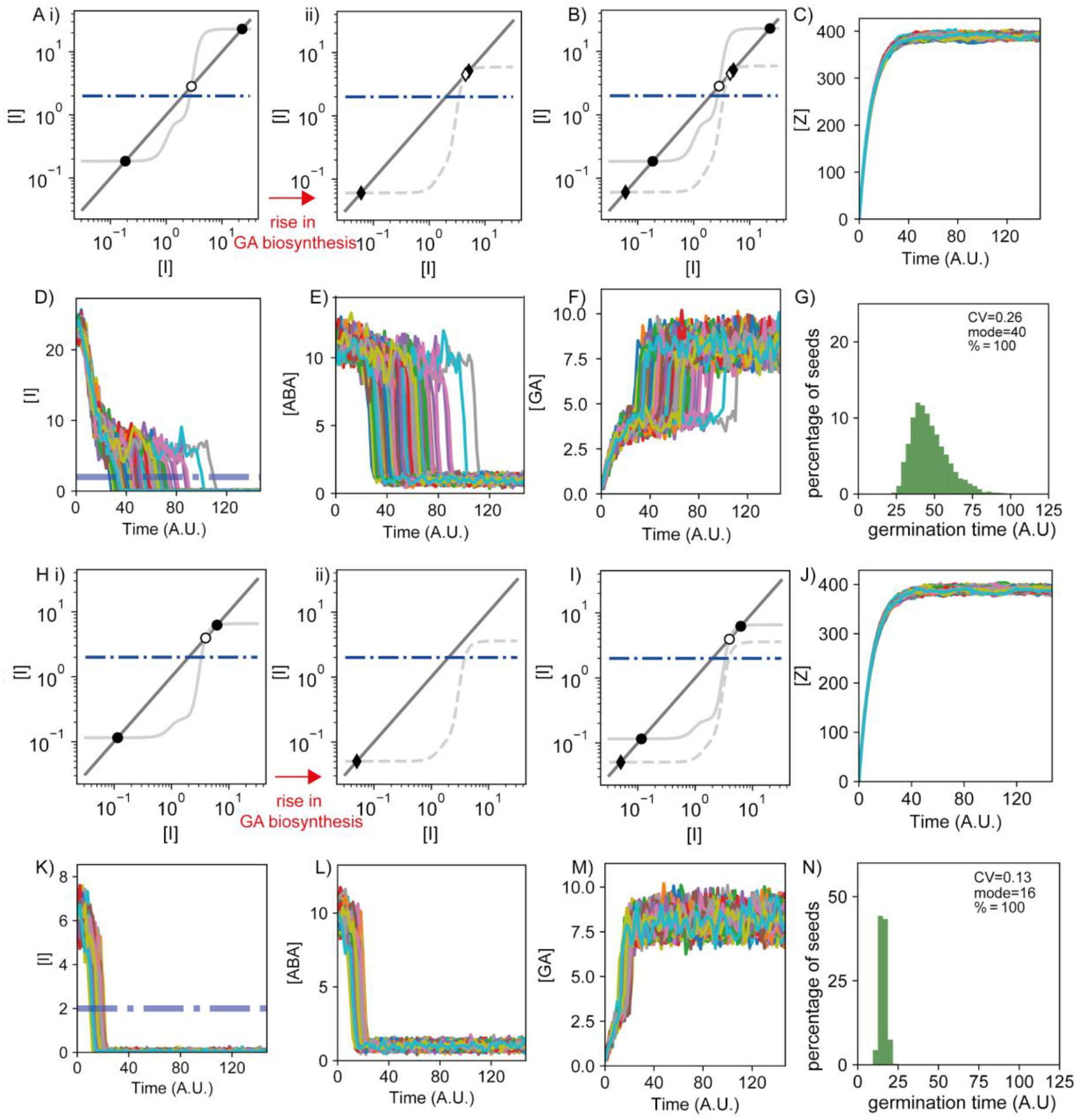
Dynamics of the components of the ABA-GA model. Modelling results showing representative behaviour of the model when it is in the bistable (A-G) and monostable (J-N) scenarios after the rise of GA biosynthesis (referred to as bistable and monostable scenarios for simplicity). The bistable scenario corresponds to the grey region in the phase diagrams shown in Figure 5-figure supplement 3, while the monostable region corresponds to the white regions. **A, H)** Results from nullcline analysis for the Integrator variable showing the steady states of the dynamics before (i) and after (ii) the GA biosynthesis increase (see methods). In each panel, steady state solutions are shown by the intersections between the dark gray line and the light grey line. Filled dots and diamonds represent the Integrator stable steady states before and after the GA biosynthesis increase, respectively. Empty dots and diamonds represent unstable steady states. The dashed-dotted blue line illustrates the Integrator threshold below which germination happens. Before the increase of GA biosynthesis, the modelled network exhibits a high Integrator stable steady state above the threshold (higher filled dot), representing a non-germinating state before sowing. We set this state as the initial condition of the simulation. For these parameter values, a lower Integrator stable solution below the germination threshold exists (lower filled dot), therefore representing a germination state, as well as an intermediate unstable Integrator solution (empty dot). Hence, bistability occurs for the Integrator variable before the increase of GA biosynthesis. With the provided noise intensity for these simulations, none of the seeds is able to switch from the non-germination state to the germination state in scenario (A), and a low percentage of seeds is able to switch in scenario (H) (see methods). A ii) In the bistable scenario after the rise in GA biosynthesis, the non-germination state (high Integrator, high ABA and low GA) approaches the unstable steady state (empty dot), becoming less stable. In this case, stochastic fluctuations enable the simulated seeds to cross the unstable steady state, reaching the germination state (low Integrator, low ABA and high GA), which becomes a more stable solution. H ii) In the monostable scenario after the rise in GA biosynthesis, the increase in GA biosynthesis has a more dramatic effect, leading the non-germination state and the unstable steady state to disappear through a saddle node bifurcation; this makes the germination state the only possible stable state. **B, I)** Nullclines analyses shown in (A) and (H) subpanels, represented together. For each panel, the light grey solid line and dots show the case before the rise in GA biosynthesis and the light grey dashed line and diamonds show the case after the rise in GA biosynthesis. **C)-F)** and **J)-M)** Time courses for the components of the model in example simulations. Different coloured lines represent different seeds. **C, J)** Time courses of the concentration of the factor Z, which increases rapidly upon sowing and promotes GA biosynthesis. **D, K)** Time courses of Integrator concentrations. Dashed-dotted blue lines show the threshold below which Integrator must drop for germination to occur. **E, L)** Time courses of ABA concentrations. **F, M)** Time courses of GA concentrations. **G, N)** Histograms of germination times, with values for CV, mode and percentage germination for the distribution. The simulation representing the bistable scenario shows a transient in which the seeds can remain in a high Integrator state until the stochastic fluctuations cause them to switch to the low Integrator state. Conversely, in the monostable scenario, the seeds achieve the low Integrator state in a more direct manner. Parameter values for (A-G) and (H-N) are the same with the exception of the Integrator degradation (*v_I_*=0.4 in (A-G) and *v_I_*=1.4 in (H-N)). See methods for further details on parameters and numerical simulations.

**Figure 5-figure supplement 2.**
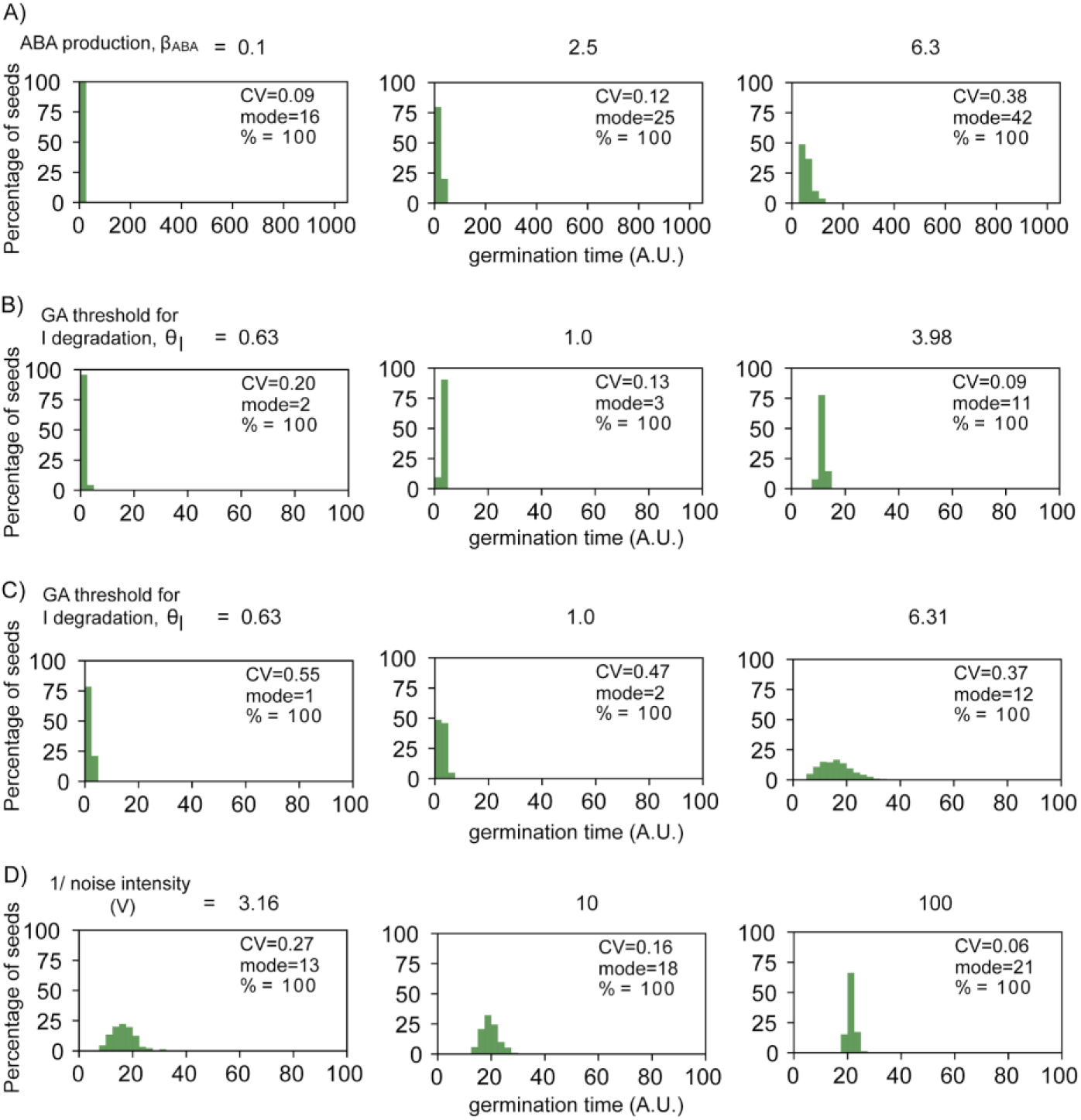
Simulated germination time distributions illustrating the effects of parameter value changes. All histograms correspond to points in the plots in Figure 5. **A)** Simulated germination time distributions for three values of basal ABA production, showing positively correlated changes in CV and mode. Data correspond to those in Figure 5B. **B)** As for A, but varying the GA threshold for Integrator degradation (which is inversely proportional to Integrator sensitivity to GA), illustrating anti-correlated changes in mode and CV. In these simulations, the ABA threshold for integrator production is set to 10. Data correspond to those in Figure 5C. **C)** As for B but varying the GA threshold for Integrator degradation in an area of parameter space where the mode increases while the CV remains relatively constant. In these simulations, the ABA threshold for integrator production is set to 6.5. Data correspond to those in Figure 5D. **D)** Varying the parameter V which governs the level of noise in the system, illustrating a change in CV while the mode remains relatively constant. Data correspond to those in Figure 5E.

**Figure 5-figure supplement 3.**
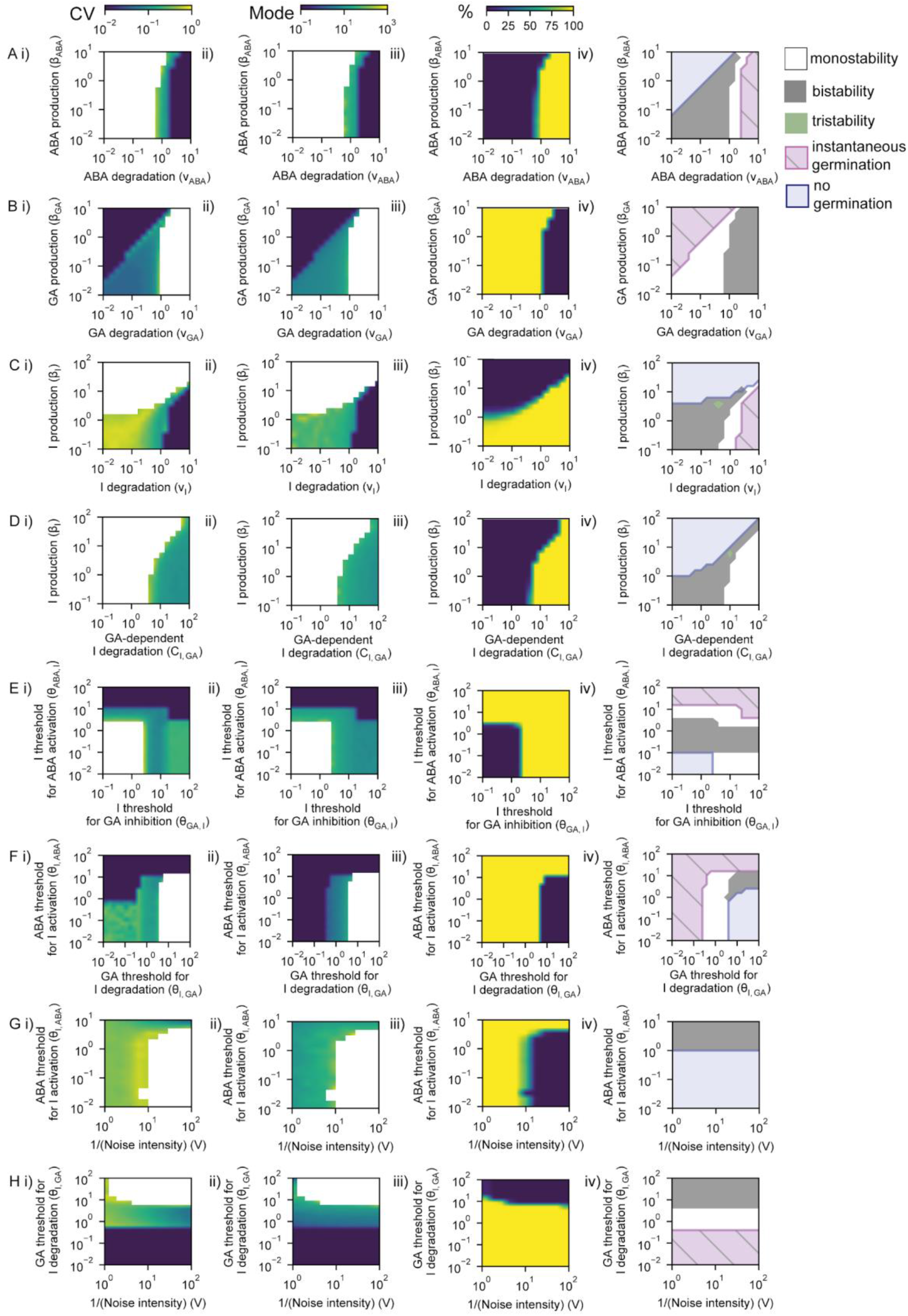
Exploring the effects of model parameters on CV, mode and percentage germination. Each panel shows a result from a 2D parameter exploration for a pair of parameters, such that each parameter is varied for a range of values of a second parameter. **A)** Effect of basal ABA production and basal ABA degradation parameters on CV (i), mode (ii) and percentage germination (%) (iii) of simulated germination distributions. CVs below 0.01 and above 1 are represented as being 0.01 and 1, respectively. CV and mode of the simulations were represented when there were more than nine seeds germinating out of 1000. iv) Phase diagram showing theoretically predicted regions from nullcline analysis of the deterministic system: bistability (grey) and tristability (green) regions in which we can expect the full range of behaviours in terms of germination, region where we expect to have all seeds germinated instantaneously (pink hatched), and region where no seeds are expected to germinate in the deterministic limit (no noise) (blue). The remaining white region is monostable and we expect all seeds (non-instantaneously) to germinate (see Figure 5-figure supplement 1 for information on this region). We expect that just the bistable and the tristable scenarios will allow a percentage of seeds to germinate that differs from 0 and 100%. The colour bars above A apply to all rows. All rows are as for A, but exploring the following parameter pairs: **B)** GA basal degradation *versus* GA basal production; **C)** Integrator basal degradation *versus* Integrator basal production; **D)** GA-dependent degradation of Integrator *versus* Integrator basal production; **E)** Threshold of Integrator for the inhibition of GA production (which is inversely correlated with sensitivity of GA to Integrator) *versus* threshold of Integrator for the promotion of ABA production (which is inversely correlated with sensitivity of ABA to Integrator); **F)** Threshold of GA for the GA-mediated degradation of Integrator (which is inversely correlated with sensitivity of Integrator to GA) *versus* threshold of ABA for the promotion of Integrator production (which is inversely correlated with sensitivity of Integrator to ABA); **G)** Effective volume of the system, V (which is inversely proportional to the noise in the system, see methods) *versus* threshold of ABA for the promotion of Integrator production; **H)** Effective volume of the system, V, *versus* threshold of GA for the degradation of Integrator. The theoretically predicted areas from nullclines (right panels) are closely predictive of the stochastic simulation outcomes (see methods). Figure 5-figure supplement 4 and Figure 5-figure supplement 5 show an analysis of the CV and mode of germination times in monostable and bistable regions of these parameter spaces. See methods for further details of the simulations and theoretical predictions and for full parameter values for each simulation.

**Figure 5-figure supplement 4.**
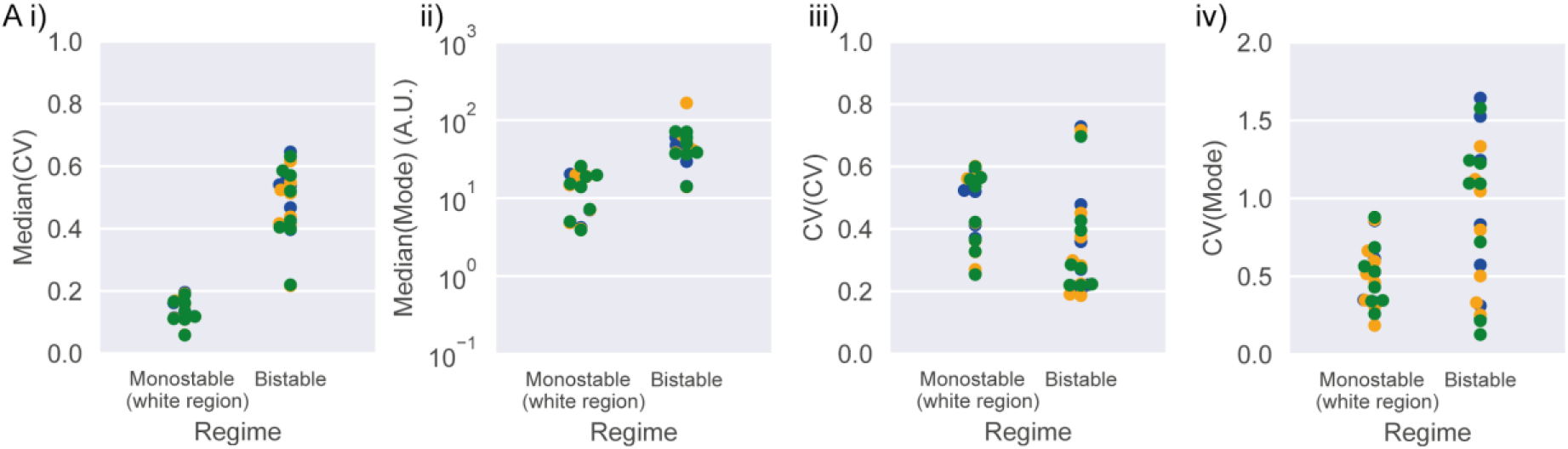
Statistics of mode and CV of germination times in bistable and monostable regions of the model after the rise in GA biosynthesis. The statistical analysis is performed on the datasets shown in Figure 5-figure supplement 5. In i) and ii) each data point is the median CV or median mode germination time, where the median is calculated across the results for a given 2D parameter screen. In iii) and iv) each data point is the CV of the CV or the CV of the mode of germination time where the CV is calculated across the results for a given 2D parameter screen. Data points falling within the instantaneous and non germination predicted regions (see Figure 5-figure supplement 3 for details of regions) were not included in this analysis because we consider them to be less biologically relevant (see methods). Colours represent different runs of stochastic simulations.

**Figure 5-figure supplement 5.**
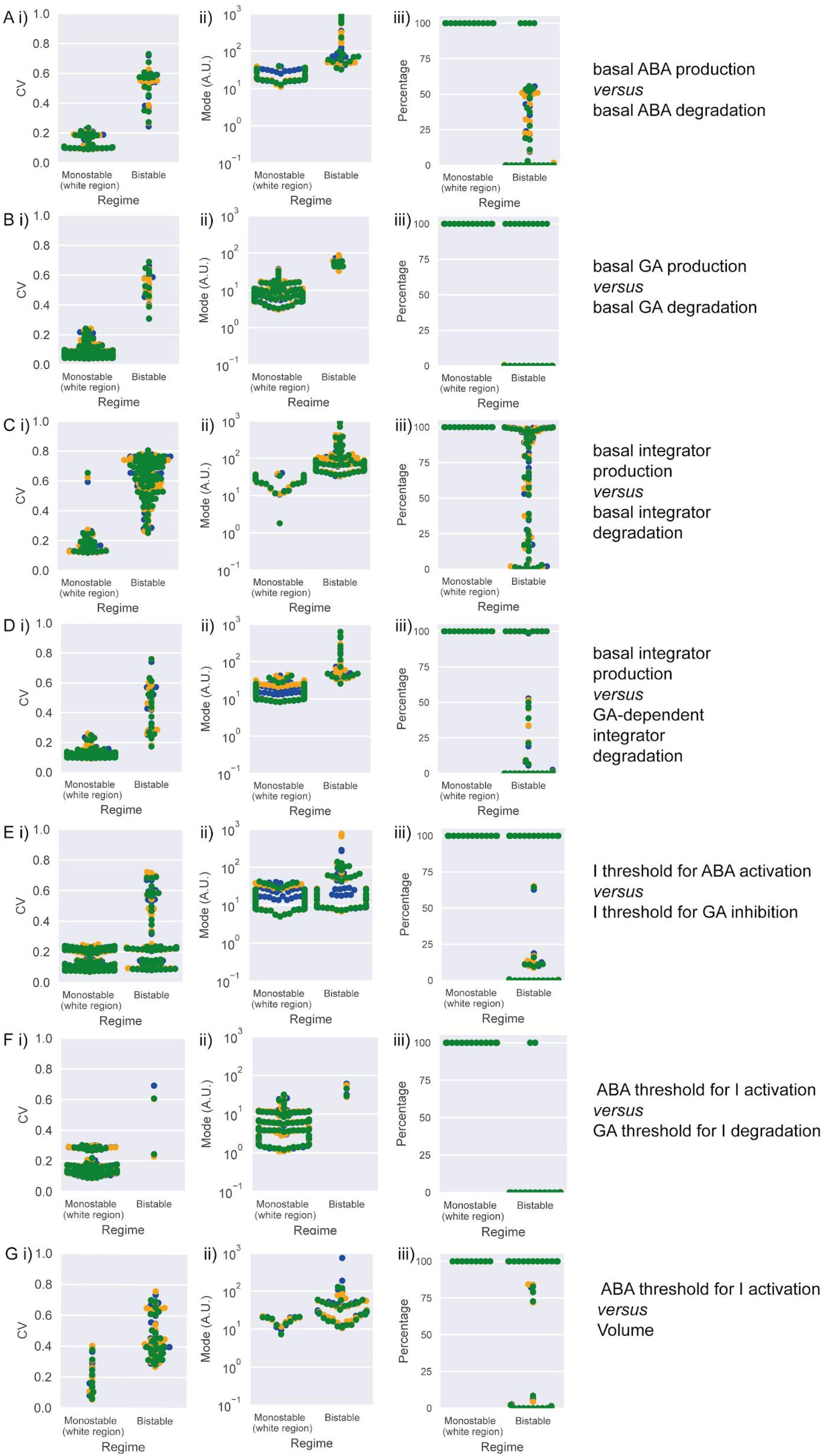
CV and mode of germination times and percentage germination in bistable and monostable regions of the model after the rise in GA biosynthesis. Simulation results across the different 2D parameter explorations shown in Figure 5-figure supplement 3. Points represent simulation results for different combinations of parameter values for the parameter pair indicated on the right. Colours represent different runs of stochastic simulations. Data points falling within the instantaneous and non germination predicted regions (see Figure 5-figure supplement 3 for details of regions) were not included in this analysis because we consider them to be less biologically relevant (see methods).

The model captures the relationships between the hormones ABA and GA and the key transcriptional regulators that act as inhibitors of germination, such as DELLAs, ABI4 and ABI5 (Ariizumi et al., 2008; Liu et al., 2016; Piskurewicz et al., 2008; Shu et al., 2016; Tyler et al., 2004). We represent these germination inhibitors as one factor, called Integrator. We model the net effects of ABA and GA on the germination inhibitors by assuming that the production of Integrator is promoted by ABA, and that its degradation is promoted by GA (Figure 5A) (Ariizumi et al., 2008; Liu et al., 2016; Piskurewicz et al., 2008; Shu et al., 2016; Tyler et al., 2004). The germination inhibitors are known to feed-back to influence GA and ABA levels through effects on their biosynthesis or catabolism (Ko et al., 2006; Oh et al., 2007; Piskurewicz et al., 2008; Shu et al., 2016, 2013). This feedback is represented in the model by assuming that Integrator promotes the production of ABA (Ko et al., 2006; Zentella et al., 2007) and inhibits the production of GA (Shu et al., 2013), (Oh et al., 2007). To capture the inhibitory effect of the DELLAs, ABI4 and ABI5 on germination, we assume that in each seed the integrator level must drop below a threshold for germination to occur. We simulate a light-induced increase in GA biosynthesis rate upon sowing (Derkx and Karssen, 1993; Oh et al., 2007, 2006). Full details and justifications of the model assumptions are provided in the materials and methods section.

The model behaves as a mutual inhibition circuit (GA inhibits Integrator and *vice-versa*) and a mutual activation circuit (ABA promotes integrator and *vice-versa*) coupled by the Integrator (Figure 5A). Overall this constitutes a double positive feedback loop that can act as a bistable switch, where there are two stable steady state solutions: high GA, low ABA and low Integrator resulting in germination; or low GA, high ABA, high Integrator resulting in no germination (Figure 5-figure supplement 1A). We hypothesised that variability in germination time is generated from stochastic fluctuations in the dynamics of the underlying gene regulatory network. To model these stochastic fluctuations, we adopt the chemical Langevin equation formalism (see methods), which takes into account the intrinsic stochasticity of the chemical reactions happening within the cell throughout time (Adalsteinsson et al., 2004; Gillespie, 2000).

Although the model is a simplified representation of the interactions between GA and ABA, it can make predictions concerning network behaviour. We found that, by modulating its parameters, we could generate a range of germination time distributions, from less variable (i.e. more peaked), to more variable (i.e. long tailed) that qualitatively match the range of germination distributions we observe experimentally (Figure 5-figure supplement 2).

We performed parameter screens in the model to investigate the extent to which particular parameters could have decoupled effects on CV and mode. Specifically, we varied the basal production and degradation rates of ABA, GA and Integrator, the parameters governing the sensitivity of the interactions between the three factors, as well as the level of noise in the system. We performed 2D parameter explorations to check the effect of varying a given parameter for a range of values of a second parameter, to ensure that the behaviours observed were robust across a range of parameter sets (Figure 5-figure supplement 3).

We found that the model could capture multiple possible relationships between the CV and mode of germination time distributions, with the nature of the relationship changing depending on which parameter was being varied. A number of parameters had positively correlated effects on CV and mode although the strength of this correlation varied depending on the region of parameter space and the parameter being changed (Figure 5B; Figure 5-figure supplement 3). We confirmed that these relationships were not due to changes in the percentage of seeds germinating (Figure 5). A positive correlation between effects on CV and mode was observed for parameters controlling all rates of basal biosynthesis and degradation (Figure 5B; Figure 5-figure supplement 2A; Figure 5-figure supplement 3A-C) as well as for those controlling the GA-dependent degradation rate of the Integrator (Figure 5-figure supplement 3D). Positive correlations between effects on CV and mode were also observed for the sensitivity of ABA production to integrator levels (we define sensitivity as the inverse of the Integrator threshold for promotion of ABA production and use the equivalent definition for all subsequent sensitivity parameters; see Figure 5-figure supplement 3E) and for the sensitivity of integrator production to the ABA level (Figure 5-figure supplement 3F). In some regions of the parameter space, anti-correlated effects on CV and mode occurred when modulating the sensitivity of the integrator to GA-promoted degradation (Figure 5C; Figure 5-figure supplement 2B; Figure 5-figure supplement 3F). In other regions of the parameter space, varying this parameter caused positively correlated changes in mode and CV with much larger changes in mode than in CV (Figure 5D; Figure 5-figure supplement 2C). Somewhat decoupled changes in CV and mode were also observed when varying the parameter that controls the level of noise in the system, such that, for some regions of parameter space, reductions in noise decreased the CV while maintaining a relatively constant mode and percentage germination (Figure 5E, Figure 5-figure supplement 2D; Figure 5-figure supplement 3G,H). Thus, the model can capture complex relationships between different germination traits.

These parameter screens also showed that across a range of parameter sets, both modes and CVs of germination times tend to be higher when bistability occurs after the rise in GA biosynthesis (Figure 5-figure supplement 3, Figure 5-figure supplement 4 and Figure 5-figure supplement 5). Additionally, the bistability region is associated with increased variability in the modal germination time between simulations (Figure 5-figure supplement 4).

To generate testable predictions, we next sought to understand how the model behaves when the levels of ABA and GA are varied through exogenous addition. The model predicts that starting from germination distributions with low or high variability, increasing concentrations of exogenous ABA will initially increase the CV of germination time distributions (Figure 6A i), causing a long-tailed distribution of germination times to emerge (Figure 6-figure supplement 1A). This is because the addition of exogenous ABA stabilises the Low GA - High ABA - High Integrator state, requiring stronger fluctuations to allow germination (Figure 6-figure supplement 2A). At higher concentrations of ABA the germination time distribution becomes flattened into a seemingly uniform distribution with a high mode and mean and therefore lower CV (Figure 6A i, iii; Figure 6-figure supplement 1A, [ABA*exo*]= 2.5). At high enough levels of exogenous ABA, the time to achieve the low integrator state becomes larger than our chosen final simulation time; seeds exhibiting this behaviour are considered non-germinating. Hence, the increase of exogenous ABA also reduces the percentage of germinated seeds (Figure 6A iv). In these cases the high exogenous ABA makes the system become monostable, causing the non-germination solution to be the only one.

**Figure 6.**
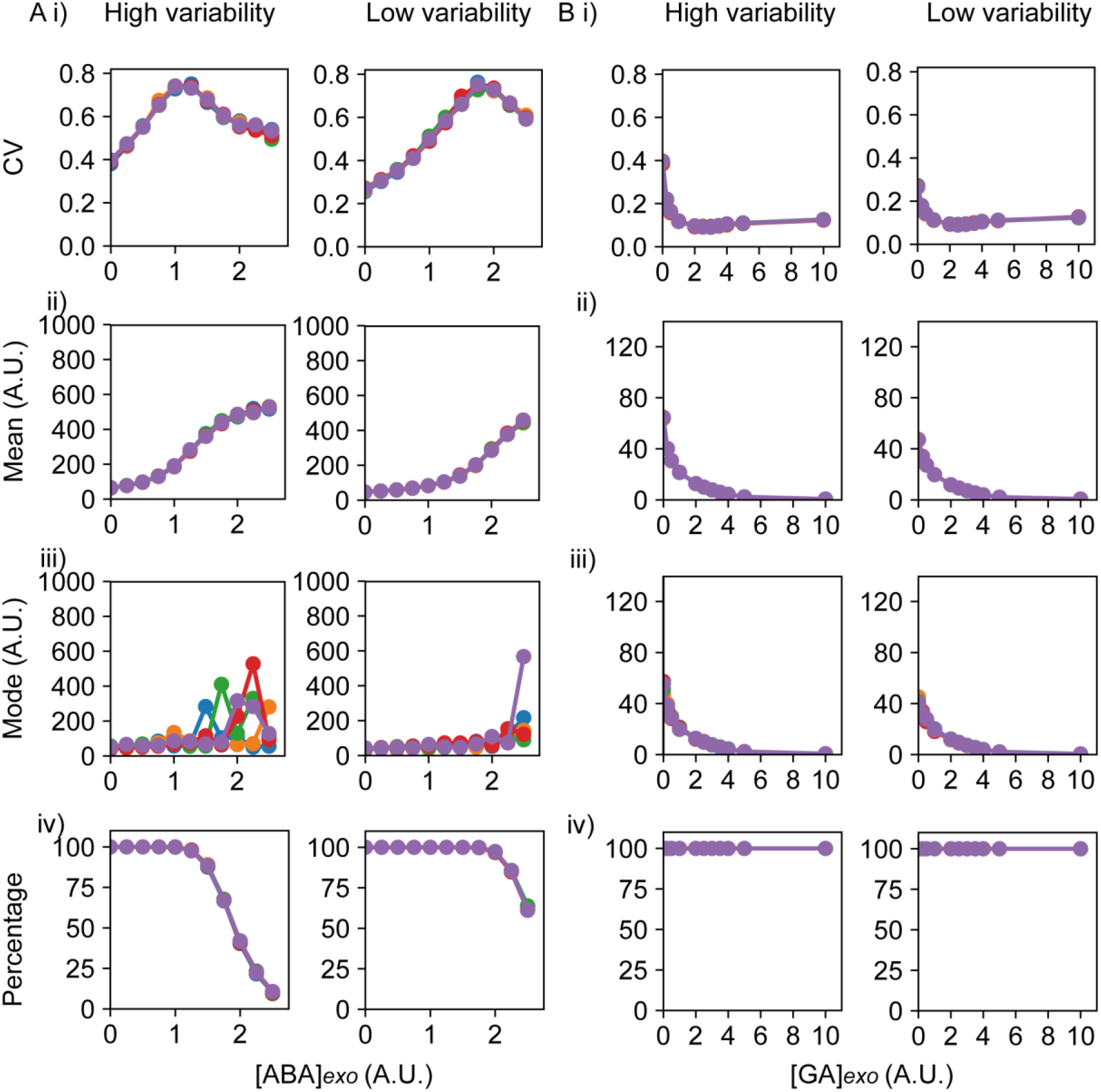
Predictions of the ABA-GA model on the effects of exogenous addition of ABA and GA. **A)** Simulations of addition of increasing doses of exogenous ABA (x axes), starting from a point in the parameter space that shows higher seed germination time variability (left) and lower variability (right) when no exogenous ABA or GA is added. Plots show the effects on the CV (i), mean (ii), mode (iii) and percentage of seeds that germinated (iv) for the resulting germination time distributions. For the ABA dose response, we chose concentrations of ABA for which there was at least 10 % germination on average for the high variability lines, to allow the other germination parameters to be ascertained. **B)** As for A, but for the addition of increasing concentrations of exogenous GA (x axes). Each panel shows the result of 5 stochastic simulations for 4000 seeds, each plotted in a different colour. Parameter values for the high and low variability lines simulations are the same with the exception of the threshold for GA degradation (θ*_I,GA_*=6 for the low variability lines and θ*_I,GA_*=6.2 for the high variability lines), and the applied hormones, which are also treated as parameters. See methods for further simulation details and parameter values. Figure 6-figure supplement 1 shows simulated germination time distributions for specific values for selected concentrations of ABA*_exo_* and GA*_exo_*. Figure 6-figure supplement 2 shows the results of nullcline analysis in the presence of ABA*_exo_* and GA*_exo_*.

**Figure 6-figure supplement 1.**
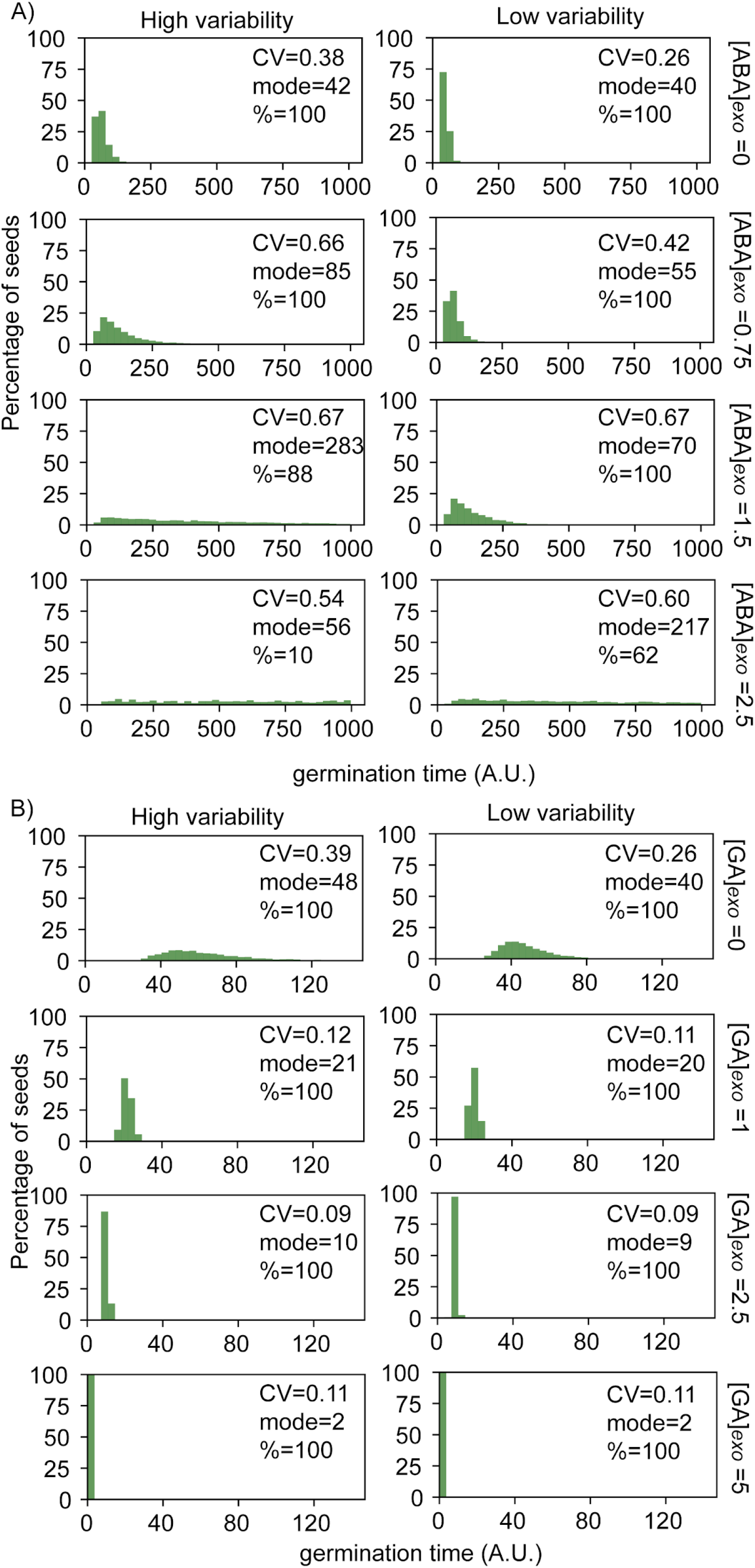
Simulated germination time distributions for a range of concentrations of exogenous ABA and GA. Simulation results of adding increasing concentrations of exogenous ABA (A) or GA (B), showing germination time distributions and the CV, mode and percentage germination for those distributions. Simulations were performed starting from a point in the parameter space that shows higher germination time variability (left) and lower germination time variability (right) when no exogenous ABA or GA is added. Results correspond to a subset of the same simulations as those presented in Figure 6. See corresponding nullclines for some of these panels in Figure 6-figure supplement 2.

**Figure 6-figure supplement 2.**
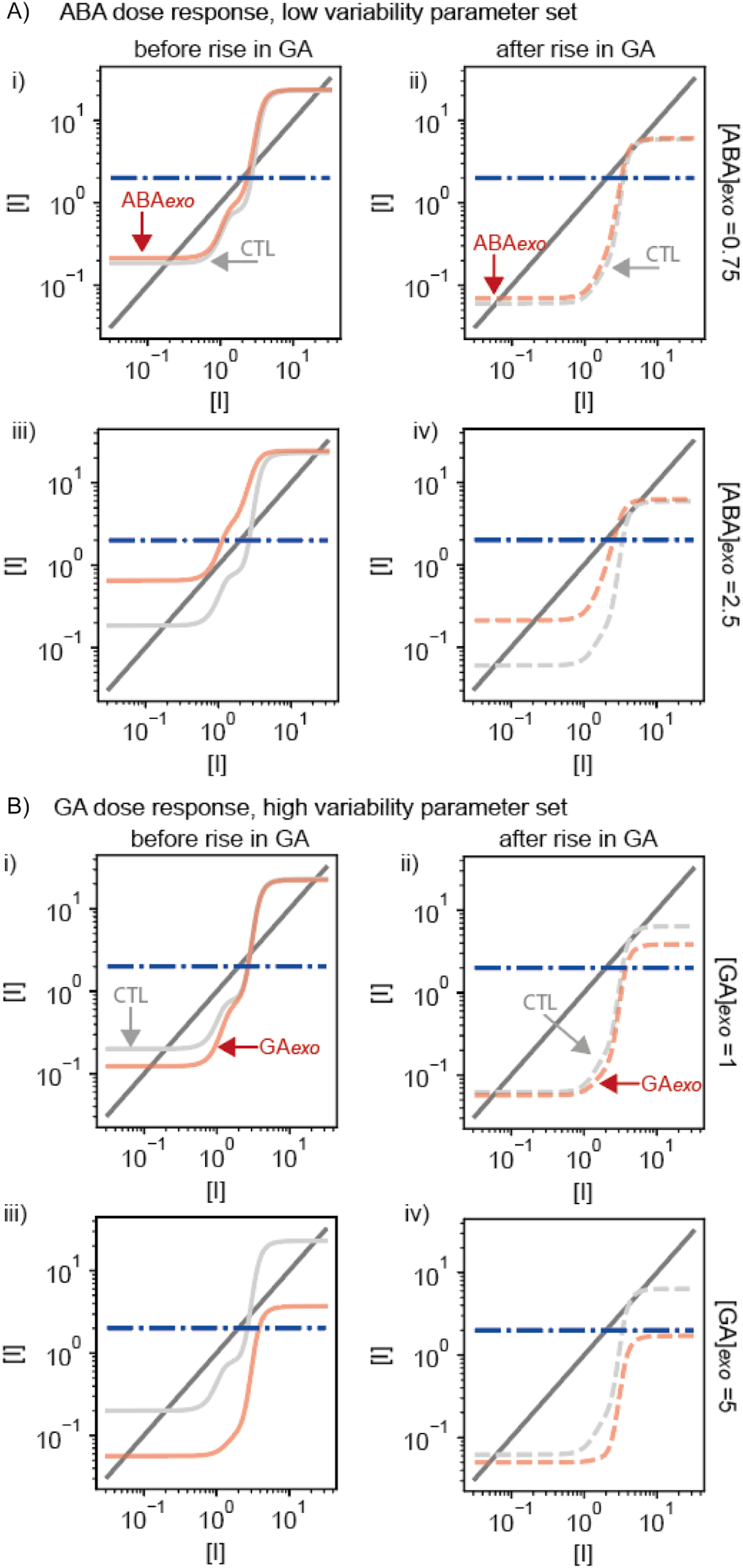
Results of nullcline analysis for ABA and GA dose responses. Plots are for example cases from Figure 6 and Figure 6-figure supplement 1. **A)** Examples of nullcline analyses for two doses of exogenous ABA applied to the low variability parameter set. Left-hand panels (i and iii) show the cases at the beginning of the simulations, prior to the rise in basal GA biosynthesis. Right hand panels (ii and iv) show plots for after the rise in basal GA biosynthesis. Light grey solid (i and iii) and dashed (ii and iv) lines show the control simulations without exogenous addition of ABA and red solid (i and iii) and dashed (ii and iv) lines show the simulations with addition of exogenous ABA (the level added is indicated on the right). Steady state solutions are shown by the intersections of the dark gray solid line with the light grey or red lines. Dashed blue line is the threshold below which Integrator must drop for germination to occur. Exogenous application of ABA (iv) can enhance the stability of the non-germination state (see methods), making it more difficult to switch to the germination state, driving a very long-tailed distribution of germination times (Figure 6-figure supplement 1A). Exogenous ABA also shifts the Low Integrator steady state higher, closer to the threshold for germination (compare red and grey lines). **B)** As for A, but for the addition of exogenous GA to the high variability parameter set. Exogenous application of GA can destroy bistability before and after the rise of GA biosynthesis, making the germination steady state the only possible steady state. This allows seeds to germinate earlier, with less variability (Figure 6-figure supplement 1B).

Conversely, addition of GA can destabilise and even destroy the Low GA - High ABA - High Integrator state (Figure 6-figure supplement 2B), readily enabling a decrease of the Integrator, and leading to a reduced mode and less variable germination times (Figure 6B i, iii; Figure 6-figure supplement 1B).

### Exogenous GA and ABA addition validates the model predictions

We next sought to test the model predictions that increasing ABA tends to increase variability in germination time, while GA decreases it. To do this we treated a number of lines with high or low variability in germination times with a range of ABA and GA doses and quantified germination at 1 day intervals.

ABA treatments tended to increase the spread of germination time distributions, particularly for low variability lines. In the low variability lines, high ABA concentrations of 5 and 10 *μ*M caused large increases in the CV of germination time, such that the distributions of germination times for low variability lines treated with ABA were similar to those of high variability lines in control conditions (Figure 7A i, Figure 7-figure supplement 1A, compare Col-0 10 *μ*M ABA with M182, 0 *μ*M ABA). This was consistent with the prediction from the model that an initial increase in ABA concentration causes an increase in CV (Figure 6A i, Figure 6-figure supplement 1A). At 5 *µ*M ABA, the increase in CV occurred with only a slight increase in mode and little change in percentage germination, whereas at 10 *µ*M there was a corresponding increase in mode days to germination and decrease in percentage germination (Figure 7A). This observation that moderate ABA levels cause a larger increase in CV compared with the change in the mode was consistent with the model (Figure 6A i, iii; Figure 6-figure supplement 1A, ABA*exo* 0 compared to ABA*exo* 1.5 (CV fold change =2.6; mode fold change=1.75)). For high variability lines, increasing concentrations of ABA increased mean and mode germination times, but had relatively little effect on variability (Figure 7A i, ii and iii, Figure 7-figure supplement 1A, 182). While the model did predict changes in CV for high variability lines, the fold-changes were smaller than those predicted for low variability lines (Figure 6A i).

**Figure 7.**
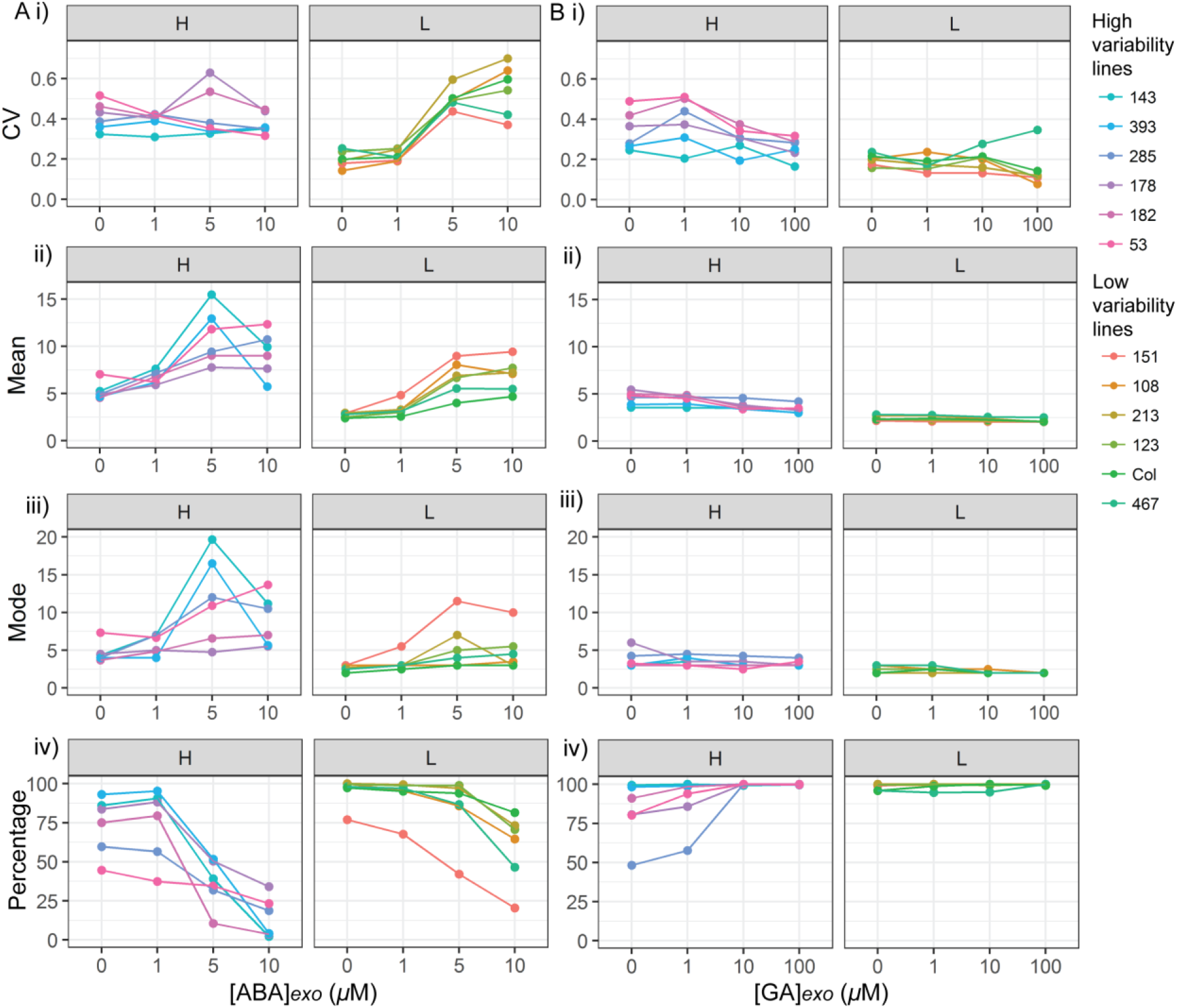
ABA and GA dose responses of high and low variability lines. **A)** ABA and **B)** GA dose responses for 6 high variability MAGIC lines (panels labelled “H”) and 6 low variability lines (5 MAGIC lines plus Col-0) (panels labelled “L”). A i) and B i) show mean CVs of individual lines for different ABA/GA concentrations (means are of at least 2 independent experiments, except for MAGIC lines 467 and 151 which have only one replicate in the GA dose response experiment), ii) mean days to germination, iii) mode days to germination, iv) percentage germination. Treatments with “0” *μ*M are vehicle control treatments. Figure 7-figure supplement 1 shows effects of ABA and GA on germination time distributions for example high and low variability lines.

**Figure 7-figure supplement 1.**
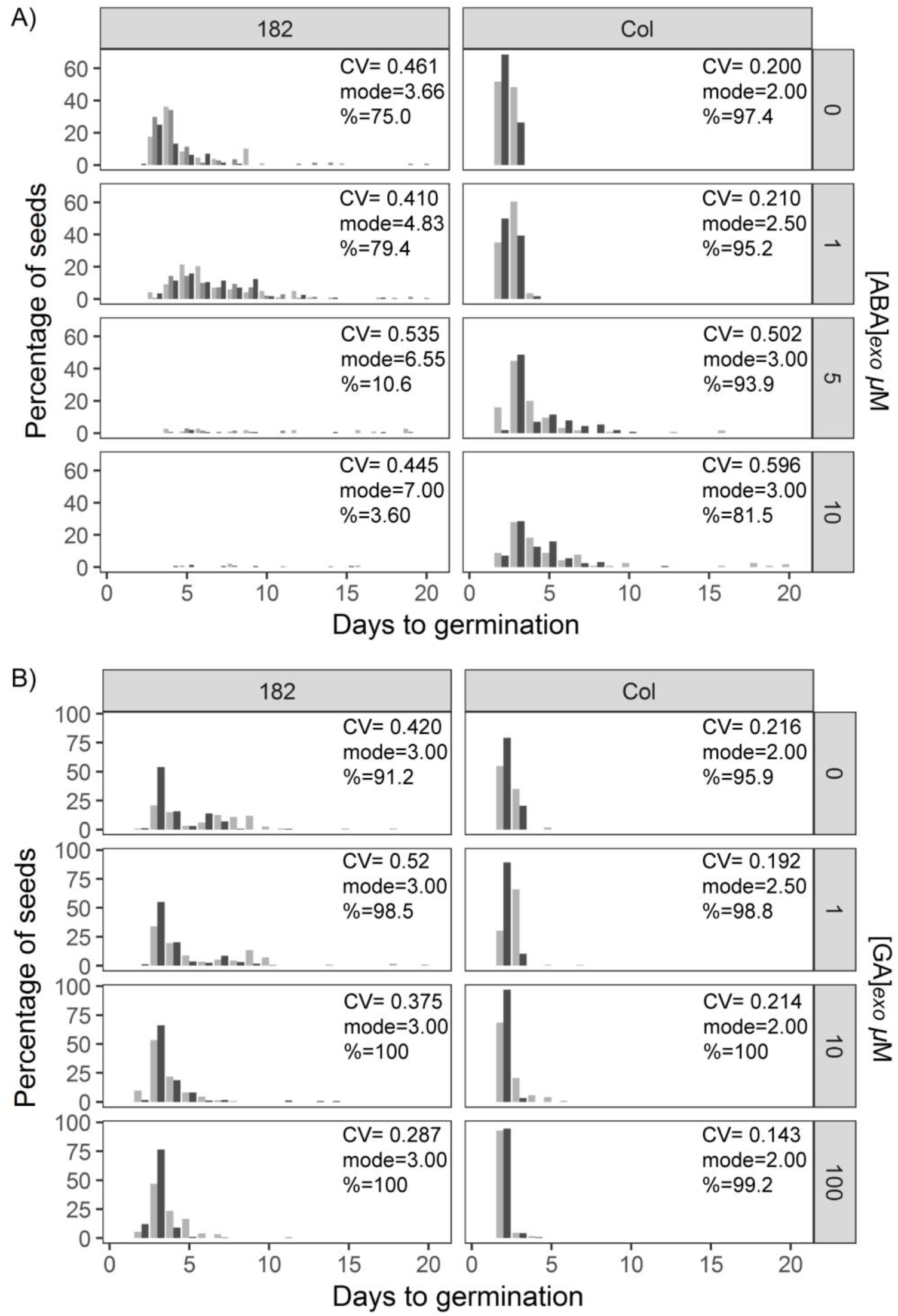
Effects of ABA and GA on germination time distributions for example high and low variability lines. **A)** Effect of increasing ABA concentration on the distribution of germination times for the high variability MAGIC line, M182 (left panels), and the low variability accession, Col-0 (right panels). Plots show the percentage of all seeds that were sown, that germinated on a given day. Horizontal rows of panels show increasing concentrations of ABA (see labels on right). Shades of grey indicate experimental replicates (at least two for each genotype). **B)** As for A, but for GA.

As predicted by the model, GA tended to decrease the level of variability in germination time, with the effect strongest for the high variability genotypes (Figure 7Bi). For high variability lines, increasing concentrations of GA tended to decrease the CV of germination time, making germination more uniform (Figure 7B, H panels; Figure 7-figure supplement 1B, 182). As expected from previous studies (Ni and Bradford 1993; Bewley 1997; Koornneef and Karssen 1994) high GA addition also increased germination percentages and decreased the mean germination time in high variability lines (Figure 7B). For low variability lines, GA had little effect on the variability (CV), percentage germination, mean or mode germination times (Figure 7B, L panels; Figure 7-figure supplement 1B, Col-0) as these lines germinated in a uniform manner with high percentage germination even in the absence of GA.

Thus, for both ABA and GA, the overall effects of exogenous addition were qualitatively similar between the model and experiments.

We also sought to investigate the effects of altered levels of ABA or GA on germination time distributions using a genetic approach. To test the effect of increased ABA concentration in the low variability background of the Col-0 accession, we used the *cyp707a1* and *cyp707a1 cyp707a2* mutants, which lack enzymes required for ABA catabolism (Kushiro et al., 2004; Okamoto et al., 2006). Similar to the effect of exogenous addition of ABA on low variability lines, loss of function of the *CYP707A1* enzyme caused the Col-0 germination time distribution to become long-tailed, this occurred without causing a large shift in the mode days to germination (Figure 8). Loss of both enzymes severely inhibited germination time and caused a large increase in the mode days to germination, similar to the effect of higher concentrations of exogenous ABA (Figure 8). This correlated effect of a change in the ABA degradation rate on CV, mode and percentage germination is consistent with the results from changing the ABA degradation parameter in the model (Figure 5-figure supplement 3A i-iii).

**Figure 8.**
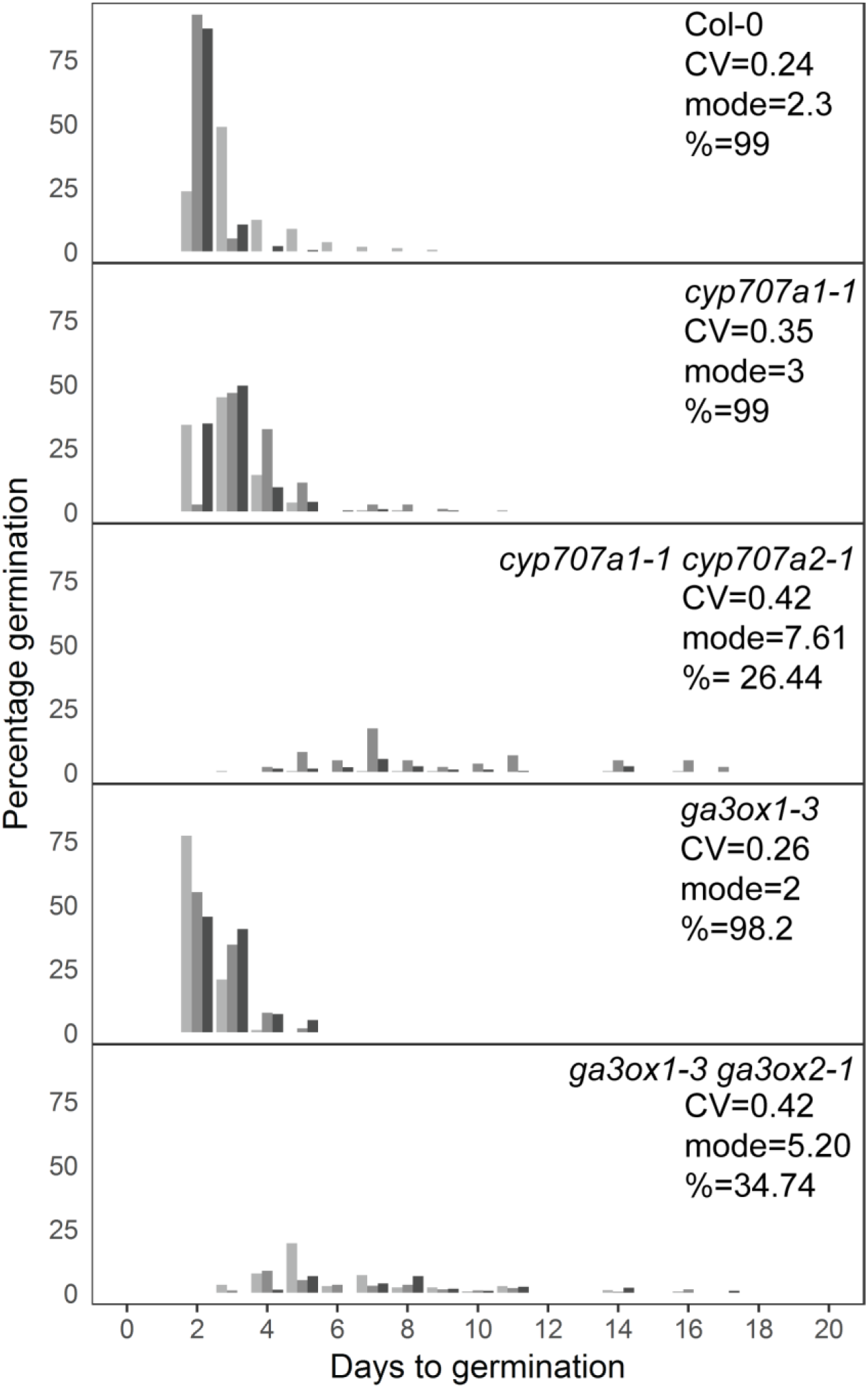
Mutants with altered levels of ABA and GA have altered germination variability. Distributions of germination times for indicated genotypes. *cyp707a1-1* and *cyp707a1-1 cyp707a2-1* mutants lack enzymes involved in ABA catabolism, while *ga3ox-3* and *ga3ox1-3 ga3ox2-1* mutants lack enzymes involved in GA biosynthesis. Grey coloured bars show the germination distribution of seed batches from replicate mother plants. For Col-0 and each mutant, the mean CV of germination times, mode days to germination and final percentage germination is shown (averaged across the replicate batches (n=3)). Data are representative of at least two independent experiments for each genotype.

To test genetically the effect of decreasing the GA concentration on the germination time distribution of Col-0, we used the *ga3ox1-3 ga3ox2*-1 mutant, which lacks two enzymes involved in GA biosynthesis (Mitchum et al., 2006). This double mutant, which has reduced GA levels (Mitchum et al., 2006) showed an increased CV and, similar to the *cyp707a1-1 cyp707a2-1* mutant, had increased mode and decreased percentage germination (Figure 8). Together with the GA and ABA dose response experiments, these findings support the model predictions regarding the effects of altering ABA and GA levels on CV, mode and percentage germination.

### High variability in germination time could function as a bet-hedging strategy

We sought to test whether high variability could be beneficial for survival in suddenly varying environments. To do this, we applied a short heat shock (49°C for 30 minutes) to high and low variability genotypes, either immediately following sowing or 5 or 8 days after sowing. This treatment kills seedlings but does not damage ungerminated seeds (Figure 9 “N” treatment (no heat shock) compared with heat shock on Day 0) (Silva-Correia et al., 2014). We scored germination each day and followed the seedlings that germinated to see if they died or survived the stress treatment. We found that higher variability lines showed a higher percentage of seedlings that survived the stress treatment when the plates were heat shocked at day 5 or day 8 (Figure 9A). This is because with low variability lines, most seeds germinated before day 5, and these seedlings then tended to be killed by the heat shock (Figure 9B, Col-0). In higher variability lines, the majority of seedlings that germinated before the heat shock were killed but late germinating seeds were still available to germinate after the stress (Figure 9B, M182). The percentage survival at the end of the experiment showed a positive correlation with both the mode days to germination and the CV of germination time for a given MAGIC line in an experiment (Figure 9-figure supplement 1).

**Figure 9.**
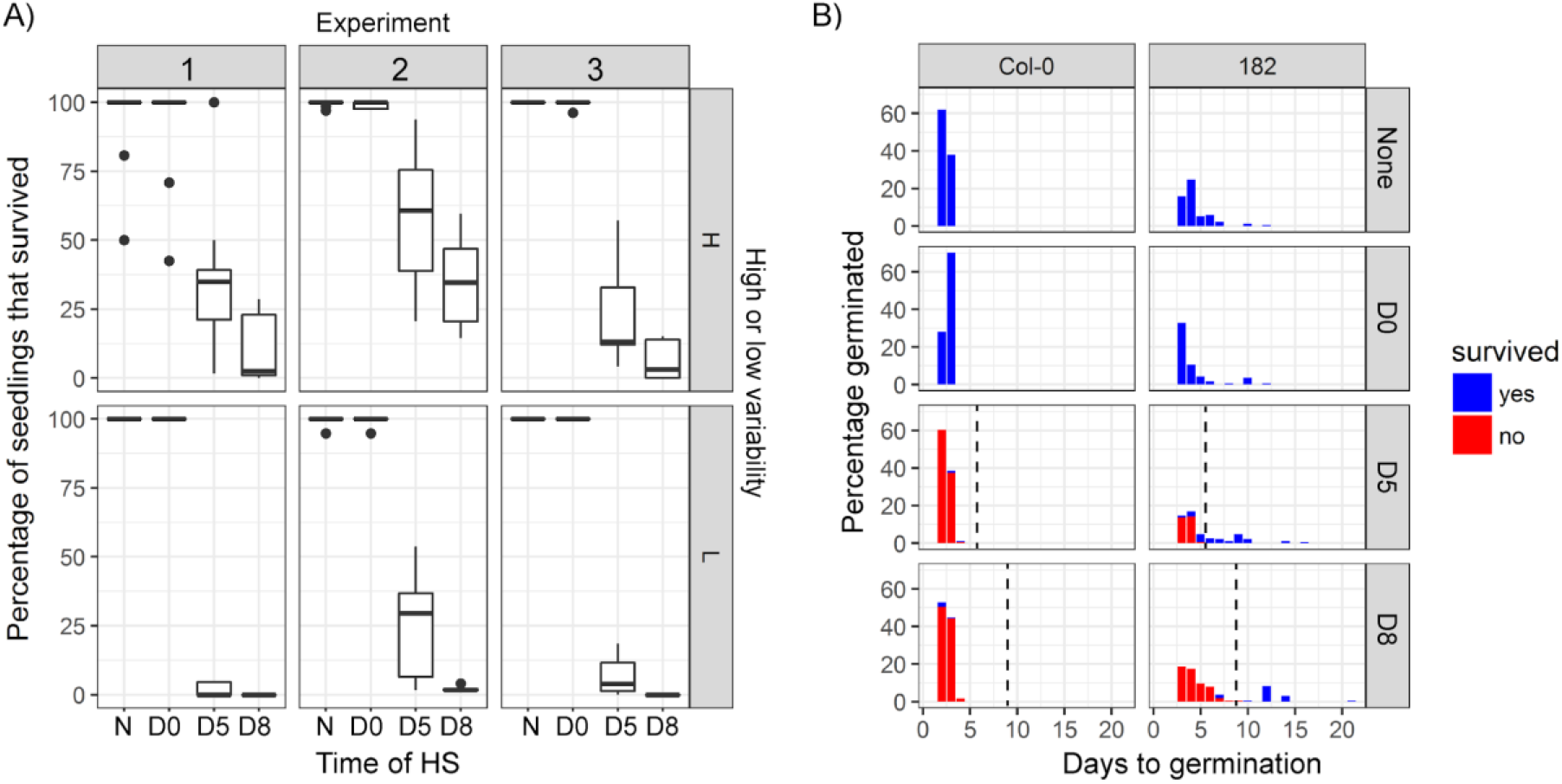
Survival of high and low variability lines when exposed to a short heat stress. **A)** High and low variability MAGIC lines were exposed to a 30 minute heat shock at ∼49°C either immediately after sowing (D0), or 5 or 8 days after sowing (D5 and D8). The treatment “N” shows non-heat shocked controls. The y axis is the percentage of seeds that germinated that survived until the end of the experiment (25 days after sowing). Experiment 1 included 3 low variability MAGIC lines, Col-0 (a low variability accession), plus 8 highly variable lines. Experiments 2 and 3 contained the lines that were present in experiment 1, plus extra lines, so that in total they included 7 low variability lines (6 low variability MAGIC lines, plus Col-0) and 9 high variability MAGIC lines. Lines were partitioned into high and low variability lines according to their average CV across the 3 experiments. **B)** Germination time distributions and survival for an example low variability line (Col-0) and high variability line (M182). Each row of panels is a different treatment. Bars show the percentage of seeds sown that germinated on a particular day. Colours of bars show the fractions of plants that germinated on a particular day that had survived or died by the end of the experiment (25 days after sowing). The vertical dashed black lines show the timings of heat shocks. Figure 9-figure supplement 1 shows how survival of a genotype relates to its mode and CV of germination time.

**Figure 9-figure supplement 1.**
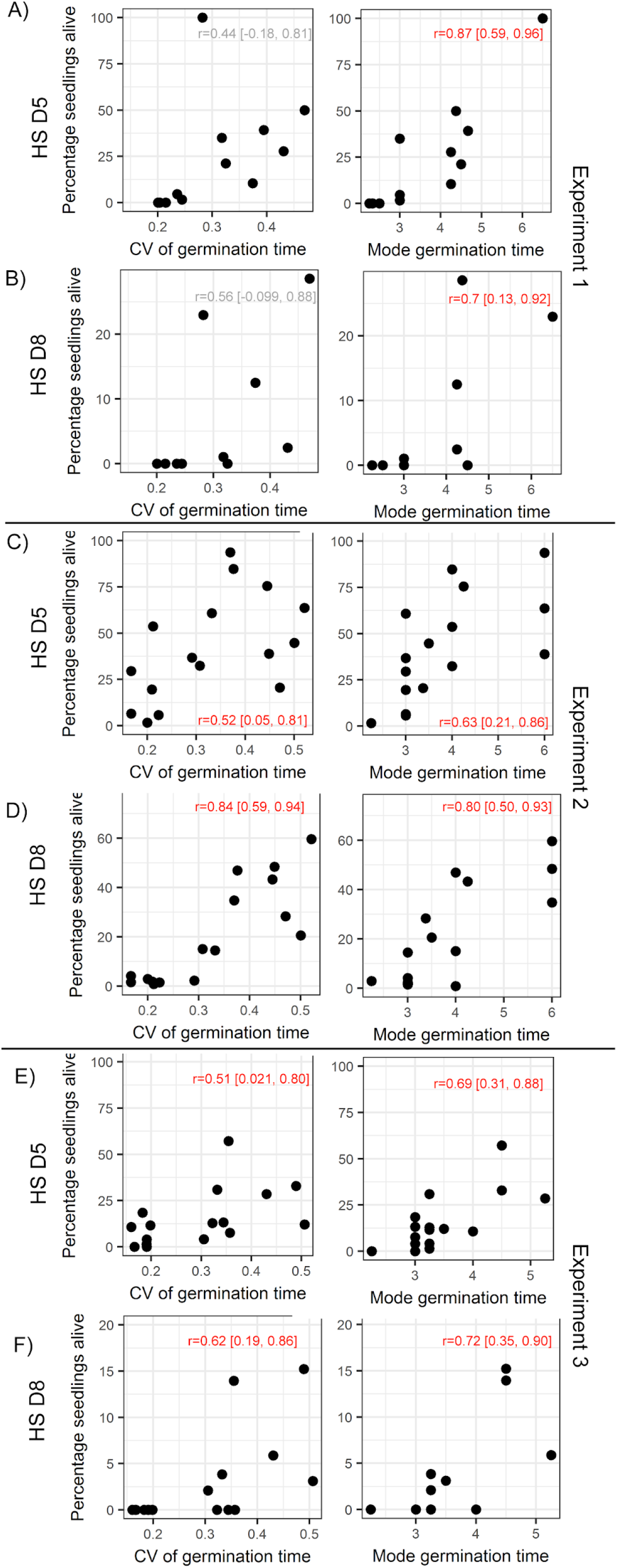
Survival of a genotype correlates with mode and CV of germination time. Scatter plots show the percentage of all germinated seedlings that survived to the end of the experiment, as a function of the CV of germination time and mode days to germination. A, C and E show results for plates heat shocked at day 5 and B, D and F show plates that were heat shocked at day 8, 3 experimental replicates are shown (same experiments as those indicated in Figure 9). On each plot, Pearson’s r is reported, along with 95% confidence intervals. Cases where there was a statistically significant correlation (with p<0.05) are shown in red.

## Discussion

Variability in seed germination time is relevant for plant survival in the wild, where high variability may function as a bet-hedging strategy. This contrasts to the situation in agriculture, where minimal levels of variability are desirable to promote crop uniformity needed for optimal harvests (Finch-Savage and Bassel, 2016; Mitchell et al., 2017). The extent to which variability in germination time is under genetic control, and therefore could be under selection, has not been well characterised. By describing detailed distributions of germination time for hundreds of Arabidopsis lines grown in a common environment, our work reveals that these distributions are genetically controlled, since they can vary greatly between different accessions and are reproducible for a given genotype. Using this natural variation, we identified two loci underlying variability, one of which affects variability relatively independently from other germination traits. Furthermore, modelling and experiments show that perturbation of the GA/ABA network can modulate variability in germination times in a predictable manner. These findings suggest that high or low variability could be specifically selected for, both in the wild, and in crop breeding programmes.

Previously it was shown that, in some Arabidopsis accessions, seeds matured on primary inflorescences had different germination behaviours to those that developed on branches (Boyd et al., 2007). It was also reported recently that some mutants defective in GA biosynthesis show graded developmental differences along the inflorescence (Plackett et al., 2018). Here we show that variability in germination time exists even for seeds from the same silique of Arabidopsis, suggesting that it is not generated by developmental gradients in the plant. We also show that seeds from proximal and distal halves of siliques have similar germination behaviours, indicating that variability is likely not caused by gradients of regulatory molecules along the length of the fruit. Thus, our findings indicate that a mechanism exists to generate different behaviours amongst seeds that are as equivalent as possible.

We show that at least two genetic loci underlie germination variability in the MAGIC population (Kover et al., 2009). The QTL at ∼16Mb on chromosome 3 overlaps with the *DELAY OF GERMINATION 6* (*DOG6*) locus (Alonso-Blanco et al., 2003; Bentsink et al., 2010), and is associated with CV, mean and mode germination time and percent germination. There is evidence to suggest that the *ANAC060* gene, which regulates ABA sensitivity, underlies this locus (Hanzi, 2014). The peak of the locus at ∼18.6Mb on chromosome 5 identified from our F2 bulked segregant mapping approach overlaps with the *DOG1* gene (Alonso-Blanco et al., 2003; Bentsink et al., 2006), which also has been shown to modulate ABA signalling (Née et al., 2017). However, the peak of the Chr 5 QTL identified in the MAGIC population lies at 19.8 Mb, equidistant between *DOG1* and a newly identified germination locus, *SET1*, which lies at around 21 Mb on chromosome 5 (Footitt et al., 2019). *SET1* was shown to influence the sensitivity of seeds buried in the soil to seasonal cycles of environmental conditions that influence germination time in the field (Footitt et al., 2019). This region contains a large number of genes involved in ABA response and seed development and it has been hypothesised that the *ABA-HYPERSENSITIVE GERMINATION1* (*AHG1*) gene, which suppresses ABA-imposed seed dormancy (Fuchs et al., 2013), underlies the effect of the locus (Footitt et al., 2019). Thus it is possible that *SET1* rather than *DOG1* underlies the Chr5 locus, or that both are relevant in the MAGIC lines, or indeed that another gene underlies the effect of this locus. Overall our findings suggest that the locus on chromosome 3 and that on chromosome 5 affect germination time variability in two different ways, with the chromosome 3 locus having correlated effects on CV, mode and percentage germination, and the chromosome 5 locus predominantly affecting the CV.

Our stochastic model of a simplified representation of the underlying GA/ABA network suggests that this bistable switch can generate variability in germination times. We show that, when in the bistable region of parameter space, the model tends to generate higher CVs and higher modal germination times compared with when operating in the monostable rapid germination region. Thus by having both possible behaviours, the model can generate a large range of “phenotypes” in terms of CV, mode and percentage germination, accounting for the variation in these traits observed in MAGIC lines and natural accessions. In line with the different effects of the genetic loci, changes to parameter values can cause a rich variety of behaviours in terms of the relationship between the CV and mode of the germination time distributions. This is consistent with the observation of a complex relationship between CV and mode in the MAGIC lines and accessions, whereby these traits are weakly positively correlated between lines, but can be partially decoupled.

Our model, which behaves as a time dependent toggle switch (Verd et al., 2014) with stochastic fluctuations, can act as a framework to understand variable germination behaviour and in future work can be used to integrate additional components of the ABA-GA network. Recent work has shown that feedback acting upon ABA biosynthesis and catabolism can modulate variability in ABA concentrations independent of the mean concentration (Johnston and Bassel, 2018). In the future it will be possible to extend our initial ABA-GA model to evaluate the effects of different feedback motifs acting on the two hormones. It is likely that with a more detailed implementation of the network, the model would have an increased ability to ascribe different germination traits and the effects of different loci to specific parameters, allowing us to make further mechanistic hypotheses about the effects of particular loci.

One limitation of our model is that we represent a seed as a single compartment in which we simulate the effects of stochastic molecular interactions. In fact the decision to germinate is likely made by groups of cells in the embryonic root (Topham et al., 2017) and thus it is unclear how the noise arising in individual cellular compartments would influence this multicellular decision. One possibility is that noise would be averaged out across the cells, reducing the level of noise at the multicellular level. This appears to be the case during vernalisation, where polycomb-based epigenetic silencing of the floral repressor *FLOWERING LOCUS C* (*FLC*) occurs in response to cold (Angel et al., 2011; Song et al., 2012). In a model of this system (Angel et al., 2015), the silencing of each *FLC* locus in each cell is proposed to be a probabilistic event, with the likelihood of silencing increasing with the duration of cold treatment. This generates heterogeneity in *FLC* expression at the individual cell level, which is then averaged out at the level of down-stream processes, resulting in the plant flowering at a time that accurately reflects the duration of cold exposure. On the other hand, noise in gene expression and phenotypic outputs has been reported for multiple pathways in plants (Jimenez-Gomez et al., 2011; Joseph et al., 2015) and recent work quantifying genome-wide transcriptional variability between individual seedlings in the same environment shows that for many genes there is a large degree of between-plant variability in transcript levels (Cortijo et al., 2019), suggesting that noise is not always averaged out to give uniformity of multicellular individuals. Rather, for some genetic networks, noise may be amplified to generate phenotypic differences. Interestingly, this transcriptomic study showed that highly variable genes were enriched for those involved in the response to ABA, supporting the idea that the bistable switch governing ABA levels amplifies noise to generate variability in gene expression and phenotypes.

Exogenous addition of GA and ABA modulates germination variability, as predicted by our model. This also fits with a population-based threshold model (Ni and Bradford, 1993) which was derived to explain the effects of GA and ABA addition on tomato seed germination distributions. In this model, the distribution of germination times for a seed batch is hypothesised to depend on the mean and standard deviation of hormone sensitivity. Similar to the effect of varying the parameters governing ABA sensitivity or production rate in our model, an increase in the mean sensitivity to a germination repressor, or an increase in the average level of the repressor, causes both an increased average germination time (increased dormancy) and a larger range of germination times (increased variability). Thus, according to this framework and consistent with the behaviour of our model, correlated changes in modal germination time, percentage germination and variability in Arabidopsis lines could be generated through variation in average sensitivities to positive or negative germination regulators, or through variation in genes that affect their average levels.

Our work reveals that the level of variability in germination times is an important trait that does not always correlate with the most common measure of germination behaviour, percentage germination. Thus to fully understand the roles of germination regulators, future work should involve characterising their effects on CV as well as on traits such as percentage germination and average germination time.

Our heat stress experiments suggest that phenotypic variability in germination time can provide an advantage, by allowing a subpopulation of seeds to survive an unpredictable period of unfavourable environmental conditions. This fits with work in desert annuals, which showed that more variable germination time distributions correlate with more variable environments (Simons, 2009; Simons and Johnston, 2006). Multiple other aspects of Arabidopsis development have been shown to display phenotypic variability, including both whole plant phenotypes such as growth rate, and molecular level phenotypes such as transcript abundance (Hall et al. 2007; Cortijo et al., 2019; Jimenez-Gomez et al., 2011; Joseph et al., 2015; Shen et al., 2012). Similar to our findings, a common feature across studies and traits is that multiple QTL tend to affect variability of a trait, and while some of them also affect the average trait value, others do not. By establishing variability in germination time as a robust trait that varies between natural accessions of Arabidopsis, our study provides the foundation for future mechanistic and functional work on phenotypic variability using this key plant trait.

## Materials and Methods

### Plant materials

MAGIC lines (Kover et al., 2009) and accessions (Vidigal et al., 2016) were obtained from the Nottingham Arabidopsis Stock Centre (NASC). The *cyp707a1-1* and *cyp707a1-1 cyp707a2-1* mutants are as described in (Okamoto et al., 2006) and were kindly provided by Eiji Nambara. *ga3ox1-3* and the *ga3ox1-3 ga3ox2-1* mutants are as described in (Mitchum et al., 2006) and were obtained from NASC.

### Phenotyping

To generate seed for assaying germination time distributions, plants were grown in batches of 40 lines in P40 trays with F2 soil treated with Intercept 70WG (both Levington, http://www.scottsprofessional.co.uk). 345 MAGIC lines and 29 accessions were phenotyped. The parental plants used for seed collection were sown in a staggered manner across 13 batches. We checked that the sowing batch of the parental plants was not a major contributor to the variation seen between lines (∼6% of the total phenotypic variance for CV could be attributed to the sowing batch, while 52% was due to the genotype of the line).

Plants for seed collection were grown in Conviron growth chambers with 16 hours of light (170 µM/m^2^/sec) and 8 hours of dark, with a day time temperature of 21°C, and night time temperature of 17°C, at 65% relative humidity. These are standard conditions for Arabidopsis growth and similar or the same as those used for seed harvest in a number of studies (Donohue et al., 2005; Finch-Savage et al., 2007; Morrison and Linder, 2014; Springthorpe and Penfield, 2015).

Some of the accessions required vernalisation to flower. For these lines, after 10 days of growth in the standard conditions described above, the plants were transferred to a Conviron growth chamber with 8 hours of light (15 µM/m^2^/sec) and 16 hours of dark, with a constant temperature of 5°C, at 90% relative humidity. For the MAGIC parental accessions that were vernalised, the plants were kept in the cold for a period of 8 weeks. For the Spanish accessions, different lengths of vernalisation period were used, as described in (Vidigal et al., 2016). Details of which accessions were vernalised, and the period of vernalisation for the Spanish accessions, are provided as Supplementary files MAGICParents.csv and SpanishAccessions.csv. To ensure that all plants used for seed harvest for QTL mapping were treated in an identical way, none of the MAGIC lines were vernalised, thus, only MAGIC lines that flowered without vernalisation were used in our study.

In a given sowing, genotypes were distributed across all trays (in random positions) and the trays were rotated approximately every 3 days to make sure that the parent plants were exposed to as similar micro-environmental conditions as possible. Six replicate plants of each line were grown. Each plant was bagged as soon as its first siliques started to ripen. Plants were watered until most (∼95%) of the siliques had ripened and then watering was stopped and plants were left in the growth chamber for 7 days to dry (Huang et al., 2014). Seeds obtained from these plants were then stored for approximately 30 days before sowing (e.g. (Morrison and Linder, 2014)), in a dark chamber kept at 15°C and 15% relative humidity. To check the quality of seed collected and stored for ∼30 days in these conditions, we performed stratification experiments for a subset of MAGIC lines (32 lines, including the most highly variable lines), by putting imbibed seeds at 4°C in the dark for 4 days prior to sowing. All but 3 lines had >90% germination after stratification, and the 3 lines that germinated poorly after stratification showed >97% germination when sown on plates containing 10*μ*M Gibberellin A_4_ (SIGMA ALDRICH, G7276).

Prior to sowing, seeds were sterilised for 4 minutes with 2.5% bleach, followed by one rinse with 70% ethanol for one minute and then washed 4 times with sterile water.

After sterilising, seeds were suspended into 0.1% agar and pipetted on to an empty petri dish, with even spacing between seeds. 0.9% agar was melted, cooled to 35 °C, then poured on top of the seeds (25 ml per round petri dish) and allowed to dry. This method of sowing seeds below agar makes scoring seeds over long time periods easier and helps to maintain a more constant environment than sowing on top of agar (where condensation forms) or on filter paper where it is difficult to maintain constant moisture levels. We checked the germination distributions for 20 lines that were sown above agar (by pipetting seeds on top of solidified and cooled agar) or below agar, as described above, and found that we obtained similar CVs of germination time for both methods. Plates were sealed with micropore tape and put into a tissue culture room with 16 hours of light (85 µM/m^2^/sec) and 8 hours of dark, a day time temperature of 20.5 °C, a night time temperature of 18.5°C, and 50% relative humidity. Each petri dish contained approximately 150 seeds. Seed germination was scored daily, using a dissecting microscope to detect radicle protrusion, and plates were checked until at least two weeks after the last germination event was observed. The germination time data is provided in data files MAGICs.csv, MAGICParents.csv and SpanishAccessions.csv. To calculate germination statistics, the data were filtered to exclude plates where less than 10 seeds germinated. We reasoned that a minimum number of seeds was needed to reliably estimate the CV. This filtering meant that 4 MAGIC lines out of the 345 that we phenotyped were excluded completely from further analysis. For 24 MAGIC lines, 1 or more replicate seed batches were excluded from further analysis. Following filtering, for 91% of MAGIC lines, at least three replicate seed batches (each collected from a different parent plant) were used for each genotype, with one petri dish for each of the three batches. For 9% of lines, only one or two replicates were used (25 lines had 2 batches, 6 had only one batch).

For 32 MAGIC lines, the whole experiment was repeated, with parental plants for seed harvest from a new independent sowing. The germination time distributions of MAGIC lines and natural accessions were all determined on agar plates as described above. The Col x No-0 F2 experiment was phenotyped on soil, in the same conditions as those described above for growing plants for seed harvest. Transparent lids were kept on the trays of soil and newly germinated seedlings were removed and counted every day. A seed was considered to have germinated when its cotyledons had visibly emerged.

### QTL mapping by bulked-segregant analysis

Col-0 x No-0 F2 seeds were sown on soil as described above. Seedlings that germinated early (day 4, early pool, E1) or in two late pools (late 1 pool, L1: days 31-39 and late 2 pool, L2: days 43 to 60) were moved into separate P40 trays and grown until flowering. The primary apices of 152 plants from E1, 321 from L1 and 213 plants from L2 were collected onto dry ice shortly after bolting and stored at −80°C. The apices from each pool were combined and ground together in liquid nitrogen and then genomic DNA was extracted using a CTAB method (Glazebrook and Weigel, 2002). Genomic DNA library preparation and sequencing was carried out by Novogene (UK) Company Limited using NEB Next® Ultra™ DNA Library Prep Kit (Cat No. E7370L). The libraries were sequenced on an Illumina NovaSeq 6000 machine with 300 cycles (150bp paired-end reads).

All of the bioinformatics processing steps and options used are detailed in the scripts provided with this paper (see data availability), so we only provide a brief summary here. We used *FastQC v0.11.3* (Andrews, n.d.) for checking read quality. Quality filtering was performed using *cutadapt 1.16* (Martin, 2011) to: remove Illumina adapters from the reads, remove reads with ambiguous base calls, and trim reads if the base quality dropped below a phred-score of 20, keeping only those reads with at least 50bp after trimming. Over 99% of bases were retained after filtering. The filtered reads were aligned to the Arabidopsis reference genome (TAIR10 version) using *bwa mem 0.7.12* (Li and Durbin, 2009) with default options, except we used the ‘`-M`’ option to mark short split alignments as secondary. Potential PCR duplicates were removed using *Picard MarkDuplicates 2.18.1* (∼20% of the reads were marked as duplicates). We performed realignment around indels using the ‘*RealignerTargetCreator*’ tool from *GATK 3.4-46* (McKenna et al., 2010). Finally, we obtained allele counts at variant sites using *freebayes v1.2.0* (Garrison and Marth, 2012) using the ‘`--pooled-continuous`’ mode, and restricting calls to sites with a depth of coverage between 10 and 400 including a minimum Phred base quality score and read mapping quality score of 20, and a minimum count of 2 and minimum frequency of 1% for the non-reference allele. We did not include indels in the analysis. To assess the presence of a QTL from these data, we used both the G’ statistic of (Magwene et al., 2011) and the simulation-based approach of (Takagi et al., 2013) to compare the three pools with each other. Both of these methods are implemented in the *R/QTLseqr v0.7.5.2* package (Mansfeld and Grumet, 2018) and gave similar results, so we report only the latter.

### QTL mapping in MAGIC lines

QTL mapping in the MAGIC lines was performed using the *happy.hbrem R* package (Kover et al., 2009) and our custom package *MagicHelpR v0.1* (available at https://github.com/tavareshugo/MagicHelpR). In summary, for each of the 1254 available markers, the probability of ancestry of an individual’s genotype at that marker was inferred using the function ‘*happy’* from the *R/happy.hbrem* package (Mott, 2015). For each marker, an N x 19 matrix is obtained with the probabilities that the N individuals inherited that piece of genome from each of the 19 founder accessions of the MAGIC population. This matrix was then used to fit a linear model, regressing the trait of interest onto this probability matrix. This type of model was used for each trait analysed. In all cases, significance was assessed using an F-test to compare the full model to a reduced model that excluded the genotype matrix, and we report the -log10(p-value) of this test.

We used a genome-wide significance threshold of -log10(p-value) = 3.5, which is an approximate threshold at α = 0.05 based on simulations (Kover et al., 2009). Significance of candidate QTL was also confirmed from a permutation-based empirical p-value based on 1000 phenotype permutations. The variance explained by candidate QTL markers was obtained from the coefficient of determination (R^2^) of the linear model.

The founder accession’s effect at each candidate QTL was estimated using the method in (Kover et al., 2009) and adapted from the *R* function ‘*imputed.one.way.anovà* in the *magic.R* script available at http://mtweb.cs.ucl.ac.uk/mus/www/magic/ (last accessed May 2020). In summary, each MAGIC line was assigned to a single founder accession based on its ancestry probabilities at that marker. MAGIC lines are then grouped by founder accession and the trait’s average is calculated for the 19 accessions. This procedure was repeated 500 times to produce an average estimate and associated 95% confidence interval (taken as the 0.025 and 0.975 quantiles of the phenotype distributions thus obtained).

All trait data were rank-transformed, to achieve normality and constant variance of the residuals in the QTL model. However, our results were robust to data transformation.

### ABA and GA experiments

Dose response experiments were performed on 6 high variability MAGIC lines (M143, M393, M285, M178, M182, M53) and 5 low variability MAGIC lines (M151, M108, M123, M213, M467), plus Col-0. Seed batches were pools of seed from 3 parent plants of each genotype. Dose responses were performed at least twice using independently collected seed batches for each MAGIC line used, except for MAGIC lines 467 and 151 which have one replicate in the GA dose response experiment.

Seeds were obtained, sowed and grown as described above for phenotyping, except that the indicated concentrations of Gibberellin A_4_ (SIGMA ALDRICH, G7276) or Abscisic acid (SIGMA ALDRICH, A1049) were added to the 0.9% agar medium used for germination assays. The GA_4_ stock was made using ethanol and the ABA stock using methanol and respective vehicle control treatments were used. Germination was scored as radicle emergence.

### Heat shock experiments

Experiments were performed for the following genotypes: low variability lines: M108, M151, M188, M200, M393, M458, M461, Col-0; high variability lines: M178, M182, M285, M304, M305, M351, M4, M492. 3 independent experiments (with seed collected from independent sowings of parent plants) were performed and gave comparable results. Seeds were obtained, sowed and grown as described above for phenotyping. In a given experimental replicate, for each stress treatment, one batch of seeds pooled from 3 parent plants was used for each line. For the heat shock treatments, petri dishes containing seeds and seedlings were wrapped in parafilm and floated in a water bath set to 49°C for 30 minutes, whilst exposed to ambient light. During the heat shock treatments there were some fluctuations of the temperature on the top of the water bath, such that temperatures ranged between 48°C and 49.5°C. Seed germination was scored every day by marking the petri dishes and 25 days after the start of the experiment petri dishes were imaged with a flatbed scanner. The images obtained were used to determine the number of seeds that germinated on each day and the number of these that survived until the end of the experiment.

### Minimal mathematical model for seed germination

We developed a model to capture the relationships between the hormones ABA and GA and key transcriptional regulators that act as inhibitors of germination. ABA and GA are known to have opposing effects on the transcription, protein levels or protein activity of the transcriptional regulators DELLAs, ABI4 and ABI5 (Ariizumi et al., 2008; Liu et al., 2016; Piskurewicz et al., 2008; Shu et al., 2016; Tyler et al., 2004). Here we represent these germinator inhibitors as one factor, called Integrator, the production of which is promoted by ABA and the degradation of which is promoted by GA.

The germination inhibitors are known to feedback to influence GA and ABA levels through effects on their biosynthesis or catabolism (Ko et al., 2006; Oh et al., 2007; Piskurewicz et al., 2008; Shu et al., 2016, 2013). For example, DELLAs promote the levels of expression of *XERICO*, which promotes ABA biosynthesis (Ko et al., 2006; Zentella et al., 2007). We capture this in the model by assuming that Integrator promotes ABA biosynthesis, creating a positive feedback loop between ABA and Integrator. Since GA inhibits the Integrator, GA ends up effectively inhibiting ABA levels through the integrator.

With regards to the feedback between the germination inhibitors (represented by the Integrator) and GA, the literature is less clear about the nature of the interaction. ABI4 appears to negatively regulate GA levels (Shu et al., 2013) supporting a double-negative (i.e. positive) feedback loop between GA levels and Integrator. However, there are mixed reports about the relationship between DELLAs and GA during germination, with studies suggesting both inhibition (Oh et al., 2007) and promotion (Topham et al., 2017; Zentella et al., 2007). As has previously been suggested (Yamaguchi and Kamiya, 2000), we assume that on balance the net relationship between the germination inhibitors and GA is negative during germination, creating a mutual inhibition between the inhibitors and GA levels. This may contribute to the large increases in GA levels that occur following sowing. With this set of interactions, since ABA increases the levels of Integrator, it effectively inhibits GA. Thus, the model captures the mutual inhibition between ABA and GA. Overall the model exhibits a mutual inhibition and mutual activation circuit coupled by Integrator (see Figure 5A), constituting a double positive feedback.

The deterministic model for ABA ([ABA]), GA ([GA] and the Integrator ([I]) is described by the following equations:

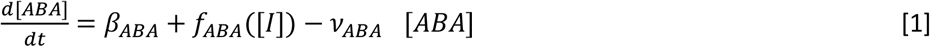

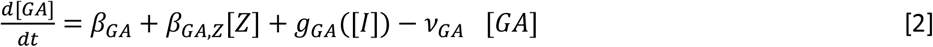

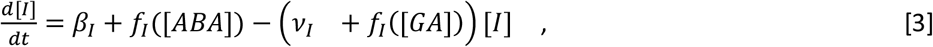

where β*_X_* and *v*_*X*_ are constitutive production and degradation rates for each *X* variable, respectively. *f*_*X*_(*y*) and *g*_*X*_(*y*)correspond to Hill increasing and decreasing regulatory functions acting on variable *X*, defined as 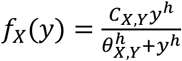 and 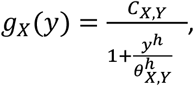 respectively, where C*_X,Y_* and θ*_X,Y_* are parameters in the function dependent on variable *Y* acting on variable *X*, and *h* is the exponent in these functions. For simplicity, we set all exponents to the same value. Note that the inverse of θ*_X,Y_* parameters can be understood as sensitivities to *Y* acting on *X*; high θ_X,Y_ values will generally require high *Y* quantities to affect *X* dynamics through the regulatory function, meaning low sensitivity of *X* to *Y*.

We simulate a sowing-induced increase in GA biosynthesis rates by adding an extra factor, [Z] which follows the dynamics governed by equation 4 and feeds into equation 2.

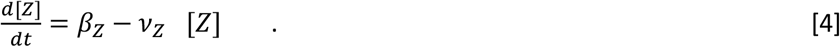

We focus on those parameters leading to either monostability, or bistability, typically showing a low GA - high ABA state and a high GA - low ABA stable state. To capture the inhibitory effect of the DELLAs, ABI4 and ABI5 (represented by Integrator) on germination, we assume that in each seed the Integrator level must drop below a threshold for germination to occur. If the system resides at the low GA state for sufficient time, the Integrator can drop below the threshold, driving germination.

Our model can be understood as a time-dependent switch (Verd et al., 2014) with stochastic fluctuations, and variability in timing -in this case, germination time - is captured when crossing a concentration threshold (Ghusinga et al., 2017). Note this circuit can also lead to tristability, but for simplicity we do not explore this model feature in detail.

In simulations of exogenous ABA or GA application, the Integrator equation follows the dynamics

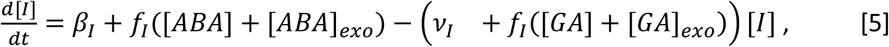

where [*ABA*]*_exo_* and [*GA*]*_exo_* are constant variables representing the concentrations of exogenous ABA and GA.

To take into account the intrinsic fluctuations of the network, we simulated the stochastic chemical Langevin equations (Gillespie, 2000); (Adalsteinsson et al., 2004) of the model equations [1-4], which read

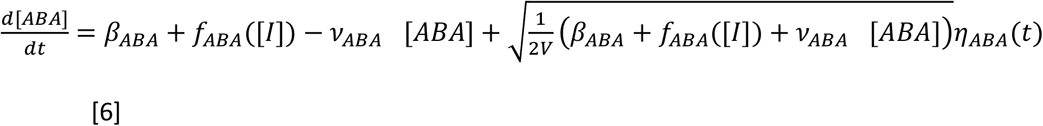

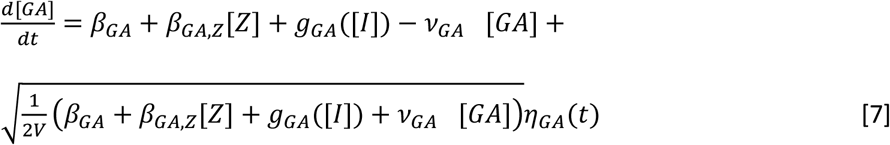

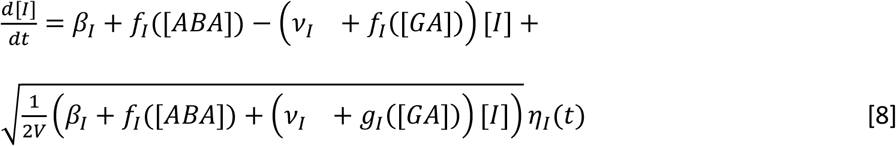

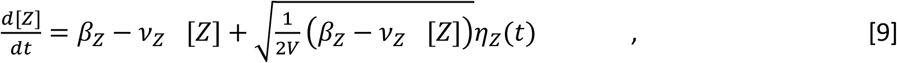

where V is an effective volume of the modelled system, which determines the strength of the stochastic term; η*_X_* is a gaussian random number with zero mean that fulfils <η*_X_*(t)η*_Y’_*(t’)>=⍰(t-t’)*⍰_X,Y_*; *⍰_X,Y_* is the Kronecker delta, where *X* and *Y* refer to concentration variables and ⍰(t-t’) is the Dirac delta, where t and t’ are two arbitrary time points. We will refer to noise intensity as the inverse of the V parameter, given that the stochastic terms are inversely proportional to V. Note that all stochastic equations recover the deterministic limit when V parameter goes to infinity, as expected for the standard chemical Langevin equation (Gillespie, 2000).

The stochastic version for equation [5] for modelling the application of exogenous ABA and GA reads

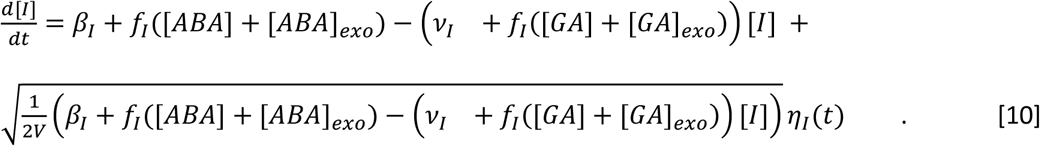

This bistable switch model is reminiscent of the bistable switch model proposed by Topham et al., 2017, although the mutual inhibition has been implemented differently, and we considered stochastic fluctuations.

Initial conditions were set at the fixed point of the deterministic model that exhibited the highest Integrator value before the sowing-induced increase in GA biosynthesis (i.e., the highest root solution for the Integrator when β_GA,Z_=0). When exogenous ABA or GA were applied, we assumed that seeds were in the same initial state as they would have been in the absence of exogenous hormone treatments. Numerical integration of the chemical Langevin equations with the îto interpretation was performed with the Heun algorithm (Carrillo et al., 2003) with an absorptive barrier at 0 to prevent negative concentration values. After each integration step, seeds were tagged as germinated if their Integrator concentration was below the germination threshold. The integration time step was set at dt=0.1. All simulations were stopped at time 1000.

Fixed points of the deterministic dynamics were computed by finding the solutions to the nullclines for the deterministic model equations [1-4], i.e., *d*[*ABA*]/*d*t=*d*[*GA*]/*d*t=*d*[I]/*d*t=*d*[*Z*]/*d*t=0, and then by substituting all the variables into the Integrator equation. The algebraic equation to solve reads

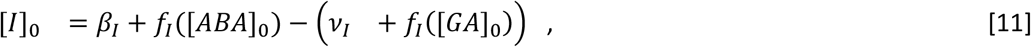

With

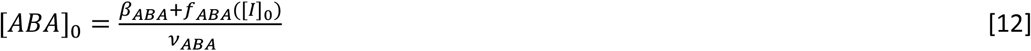

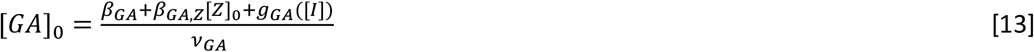

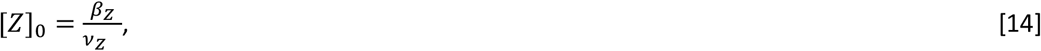

where [*ABA*]_0_, [*GA*]_0_, [*I*]_0_ and [*Z*]_0_ are the steady state solutions of the different variables. For finding the fixed points in the cases of exogenous application of GA and ABA, an equivalent procedure was performed with Equations [1-3] and [5].

To find all the solutions at each particular parameter set, we first used the bisection method throughout logarithmically spaced intervals for the Integrator variable to find approximate solutions, and then we used the opt.brentq scipy function in Python to find the exact solutions. We also represented the left and right hand side of equation [11] or an equivalent equation for the exogenous application of GA and ABA to graphically see the solutions of the system (see Figure 5-figure supplement 1 and Figure 6-figure supplement 2). In these plots, we represented the deterministic stability by analysing the sign of dI/dt at the vicinity of the solutions (Strogatz, 2015). Upon the variation of a certain parameter value or the application of exogenous ABA or GA, a given stable fixed point will approach to (or get further from) the separatrix, the hyperplane in the solution space separating the basins of attraction of the stable fixed points (Strogatz, 2015), which contains an unstable fixed point in our case. When studying the stochastic system, we will assume that a stable fixed point will lose stability when it approaches the unstable fixed point, given that this most likely will facilitate the stochastic switching to the other stable fixed point; conversely, a stable fixed point will gain stability when it gets further from the unstable fixed point. This assumption is consistent with the outcome of our simulations (Figure 6-figure supplement 1 and Figure 6-figure supplement 2).

Unless otherwise stated, parameter values were set to the following values (all units are arbitrary): β*_ABA_*=1, β*_GA_*=0.3, β*_GA,Z_*=0.01, β*_Z_*=39 and β*_I_*=0.3 for the production rates; *⍰_ABA_*=*⍰_GA_*=1, *⍰_Z_*=0.1 and *⍰_I_*=0.4 for the degradation rates; thresholds were θ*_ABA,I_*=3.7, θ*_GA,I_*=1.2, θ*_I,ABA_*=6.5 and θ*_I,GA_*=6; coefficients for the regulatory functions are C*_ABA,I_*=10, C*_GA,I_*=4, C*_I,ABA_*=10 and C*_I,GA_*=6; exponents in the regulatory functions were all set to *h*=4; the effective volume was set to V=30; the Integrator threshold for germination was set to 2. For Figure 5, in B, *⍰_ABA_* =1.58; in C and E, θ*_I,ABA_*=10; in D, V=1.

To better understand the dynamics across several parameter ranges, we performed simulations varying 2 parameters at the same time. The resolution of the parameter exploration was of 4 to 5 parameter values per order of magnitude, logarithmically spaced. In this parameter space exploration, we also studied the different regions of the parameter space that could be predicted from nullcline analysis. This allowed us to find the bistable and tristable regions of the parameter space, the regions where no germination is expected in the deterministic limit and the regions where germination would instantaneously occur. The remaining regions in the parameter space were monostable. Note that monostable, bistable and tristable regions were computed by counting the number of steady states after the rise of GA biosynthesis. Regions where no germination would occur in the deterministic limit were those where the lowest fixed point for the integrator was higher than the germination threshold. Regions where germination would instantaneously occur were those regions having the highest Integrator fixed point below the germination threshold. CV and mode of the simulations were represented when there were more than nine seeds germinating out of 1000, so the percentage of germination was equal to or higher than 1 %.

The theoretical regions across the parameter spaces closely predicted through nullcline analysis of the deterministic model the stochastic simulation outcomes (Figure 5-figure supplement 3). One exception was that simulations showed that occasionally some germination happened after a small number of simulation steps in the instantaneous germination region. In these cases, even if initial conditions were below the germination threshold, after an integrator step, the Integrator variable was not below the threshold anymore due to the stochastic fluctuations, and therefore, germination happened later on during the simulation (Figure 5-figure supplement 3F). Also, in the area where no germination was expected in the deterministic limit, germination could occur in some occasions due to stochastic fluctuations, leading to different germination percentages. This happened either when the germination threshold was just below the lowest target fixed point after the rise of GA biosynthesis, or at high noise intensities (e.g., see Figure 5-figure supplement 3E, G).

Our theoretical analysis and simulations showed that there are two different prototypical dynamical behaviours that are most biologically relevant (Figure 5-figure supplement 1). On one side, simulations in which there is bistability after the rise of GA biosynthesis, where seeds can undergo a transient in which they remain in a high Integrator state above the germination threshold, until stochastic fluctuations make them switch to the low Integrator state, driving germination. On the other side, simulations in which there is monostability after the rise of GA biosynthesis, where the seeds achieve the low Integrator state in a more direct manner. Those simulations falling within the instantaneous germination region would not be biologically relevant, given that a certain time is needed for seeds to germinate after sowing. Simulations leading to germination and falling within the non germination region in the deterministic limit would also be less biologically relevant, given that the Integrator would repeatedly cross the threshold back and forth, not persisting below it.

For simplicity, throughout the text, we call monostable regions those regions in the parameter space that are monostable after the rise of GA biosynthesis and are biologically relevant (i.e., the white theoretical predicted regions in Figure 5-figure supplement 3). Note the instantaneous germination regions and the no germination regions can also contain monostable, bistable and tristable cases. In Figure 5-figure supplement 4 and Figure 5-figure supplement 5 we exclude areas of parameter space that have these less biologically relevant behaviours.

The parameter space explorations and derived panels show stochastic simulation runs for 400 seeds (Figure 5; Figure 5-figure supplements 2-5). The dose dependence plots and derived panels show simulation runs for 4000 seeds (Figure 6; Figure 6-figure supplement 1), as well as simulations shown in Figure 5-figure supplement 1. Two additional stochastic simulations on 40 seeds were run in relation to Figure 5-figure supplement 1A, H, to study whether germination occurred before the rise of GA biosynthesis (i.e., for β*_GA,Z_*=0). We found that no germination occurred for the scenario shown in panel (A), while a low percentage of seeds germinated (12.5 %) for the scenario shown in panel (H).

Numerical simulations were performed with the Organism simulator (https://gitlab.com/slcu/teamHJ/Organism (Jönsson et al., 2005). Modelling figures were produced with the Matplotlib Python library (Hunter, 2007).

## Data availability statement

Whole genome sequence data was deposited to NCBI’s Short Read Archive (BioProject accession PRJNA486286). All data analysis and modelling scripts can be found at https://gitlab.com/slcu/teamJL/abley_formosa_etal_2020. Both the raw and processed experimental data for use with the analysis scripts are available from the Cambridge Apollo Repository (doi:X - persistent DOI will be available upon acceptance, but reviewers can provisionally download the data from: https://drive.google.com/file/d/1-3YKuCFcRYVk-B4RW1ZBZ6GkplVHCM3D/view?usp=sharing).

## Acknowledgements

We thank Sandra Cortijo for critical reading of the manuscript and Mana Afsharinafar, Casandra Villava, Ting Wang and Helena Kelly for help with taking care of plants and seed scoring. P. F.-J. thanks Ruben Perez-Carrasco for fruitful discussions about the modelling. Work in the Locke and Leyser labs was supported by fellowships from the Gatsby Charitable Foundation (Locke lab: GAT3272/GLC and Leyser Lab: GAT3272C).

